# Organ protection by caloric restriction depends on activation of the *de novo* NAD+ synthesis pathway

**DOI:** 10.1101/2021.08.20.457042

**Authors:** Martin Richard Späth, Karla Johanna Ruth Hoyer-Allo, Marc Johnsen, Martin Höhne, Christina Lucas, Susanne Brodesser, Jan-Wilm Lackmann, Katharina Kiefer, Felix Carlo Koehler, Petra Schiller, Torsten Kubacki, Franziska Grundmann, Thomas Benzing, Bernhard Schermer, Volker Burst, Roman-Ulrich Müller

## Abstract

Therapeutic strategies to treat acute kidney injury (AKI) are lacking in clinical practice. Interestingly, preconditioning by hypoxia (HP) and caloric restriction (CR) is highly protective in rodent AKI models. However, the underlying molecular mechanisms of this process are unknown. A comparative transcriptome analysis of murine kidneys after HP and CR identified Kynureninase (KYNU) as a common downstream target. Using a newly generated KYNU-deficient mouse line, we show that KYNU strongly contributes to the protective effect of preconditioning. Metabolome, transcriptome and proteome analyses reveal the KYNU-dependent *de novo* nicotinamide adenine dinucleotide (NAD+) biosynthesis pathway as necessary for CR-associated maintenance of NAD+ levels. Importantly, the impact of CR on the *de novo* NAD+ biosynthesis pathway can be recapitulated in humans. These findings provide a valuable insight into the molecular mechanisms mediating protection upon preconditioning and point towards the *de novo* branch of NAD+ biosynthesis as a conserved target in nephroprotection.

## Introduction

Acute kidney injury (AKI) is a frequent complication in hospitalized patients (1) with an increasing incidence over the last decades (2). Even mild forms of AKI are strongly associated with increased morbidity and mortality (3–8).

Besides sepsis and exposure to nephrotoxic agents, ischemia-reperfusion injury (IRI) is one of the main causes of AKI, which may occur as, e.g. a consequence of major surgery (9, 10). Patients suffering from chronic kidney disease (CKD), arterial hypertension, diabetes and heart disease are at increased risk of developing AKI (11). Unfortunately, targeted measures to treat or even prevent AKI are as yet lacking, making maintenance of euvolemia the current mainstay in patients at risk (9, 12).

In rodent models of AKI, preconditioning strategies employing non-harmful levels of defined stressors have shown a robust and highly potent protective effect (13–17). Currently, preconditioning by hypoxia (HP) and caloric restriction (CR) are among the most efficient approaches (14, 18–20). However, translation to the human setting is hampered by limited feasibility in the heterogeneous cohorts of multimorbid patients in the clinical setting. Thus, basic research aimed at unraveling the molecular mechanisms that underlie the protective potential of preconditioning is urgently required to refine and expand on these limited therapeutic preconditioning strategies and identify druggable targets.

In a recent publication (18), we hypothesized that HP and CR may modulate similar downstream targets in transcriptome and metabolism. This study identified Kynureninase (KYNU) – a central enzyme in the tryptophan metabolic pathway– as one of the top candidates induced by both preconditioning strategies. The tryptophan metabolic pathway is the basis of *de novo* nicotinamide adenine dinucleotide (NAD+) biosynthesis (21), which has gained increasing attention over the last years due to its protective potential in IRI (20). Based on these findings and using a novel knockout mouse line in combination with metabolome, transcriptome and proteome analyses, this study examines the contribution of KYNU to renal organ protection.

## Results

### Preconditioning-mediated induction of *Kynu*

In a recently published study we had used the comparison of two modes of preconditioning – HP and CR – to examine the underlying mechanisms of renal organ protection (18). Male C57Bl/6J mice were divided into three treatment groups (Fig. 1A). The first group underwent IRI without preconditioning serving as a control group (nonPC). The second group was exposed to 3 days of transient HP and the third group underwent 4 weeks of 66 % CR. On the day of surgery, all mice underwent a right nephrectomy followed by unilateral warm ischemia of the left kidney for 40 minutes. To obtain blood and the left damaged kidney, all mice were sacrificed 24 h after reperfusion (Fig. 1A).

**Fig. 1:**
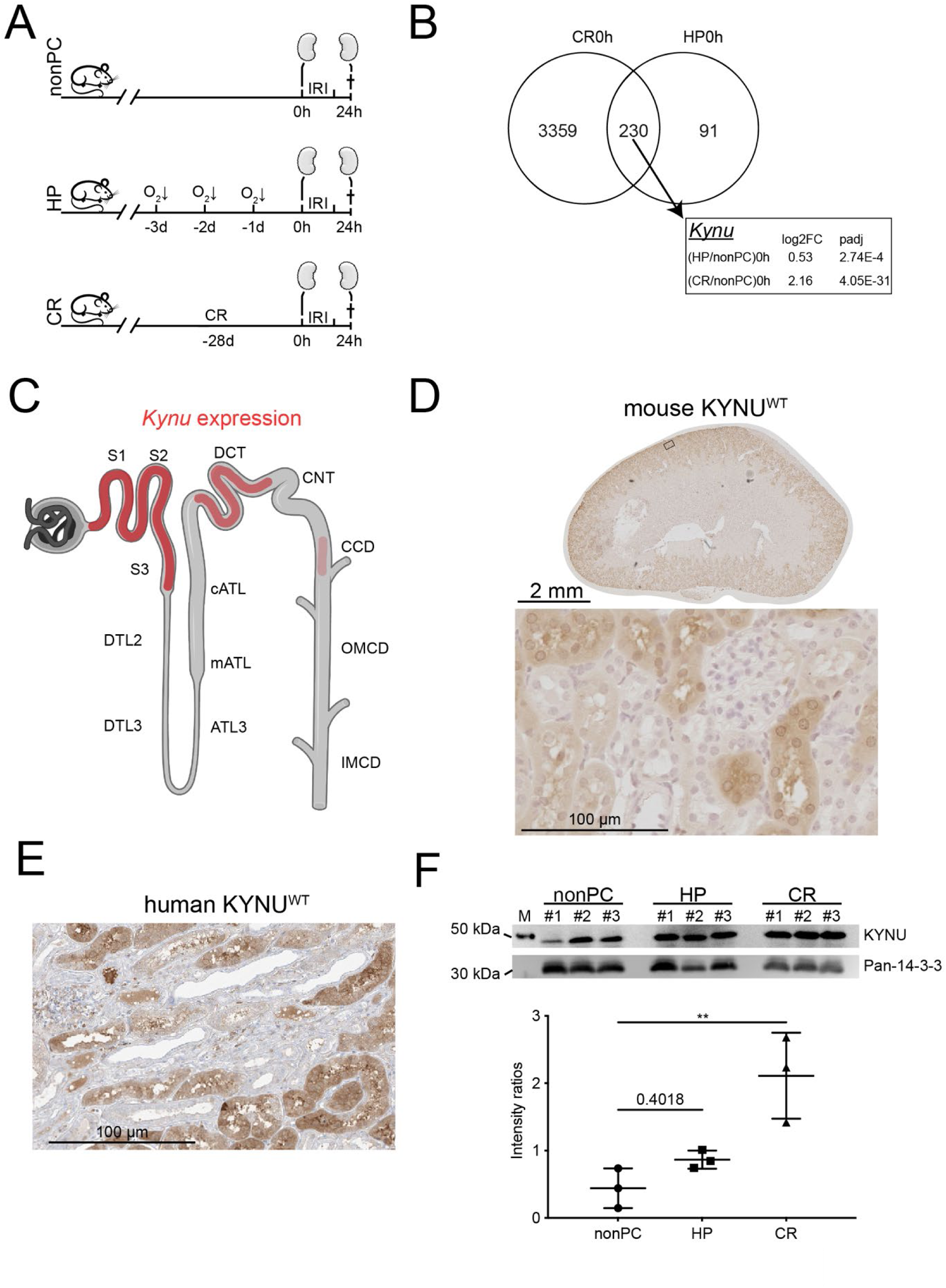
Kynureninase is expressed pre-dominantly in the proximal tubulus and induced by two modes of preconditioning. **A.** Schematic illustration of the experimental setup modified from Johnsen et al. JASN 2020 (18). C57Bl/6J mice were divided into three treatment groups (nonPC, HP, CR) prior to IRI. Right nephrectomy was performed at the timepoint of surgery (0 h). To obtain blood and the left kidneys, 24 h after reperfusion all mice were sacrificed. **B.** Venn diagram illustrating genes only regulated by CR (3359), HP (91) or both (230) at 0 h prior to IRI modified from Johnsen et al. JASN 2020 (shiny.cecad.uni-koeln.de:3838/IRaP/ (18)). Preconditioning-induced expression changes of *Kynu* pre-IRI (CR vs. nonPC and HP vs. nonPC) are displayed in the box. **C.** Schematic illustration of the segmental localization of *Kynu* along the nephron compiled from previously published single-cell (22), single-nucleus (23) and bulk-RNAseq (24) data as well as proteome analyses of microdissected kidneys (25). **D.** Immunohistochemical staining of KYNU in murine kidneys confirms the predominant localization in the proximal tubulus. **E.** Immunohistochemical staining of an adult human kidney using anti-KYNU antibody (HPA031686, derived from www.humanproteinatlas.org (26)). **F.** Immunoblot analyses of murine kidneys showing the expression of KYNU in non-preconditioned mice as well as after HP and CR (loading control: Pan-14-3-3). Intensity ratio analysis by one-way ANOVA and Tukey posthoc test reveals a significant induction of KYNU on the protein level by CR (** = p<0.01) whereas the induction by HP did not reach statistical significance (p-value>0.4018). **Abb.:** nonPC: non-preconditioned male mice; HP: hypoxic preconditioning; CR: caloric restriction; d: day; IRI: ischemia-reperfusion injury; log2FC: log2Foldchange; padj: adjusted p-value; S1 Proximal tubule; S2 Proximal tubule; S3 Proximal tubule; descending thin limb of Henle’s loop, short-loop; descending thin limb of Henle’s loop, long-loop, outer medulla; descending thin limb of Henle’s loop, long-loop, inner medulla; ascending thin limb of Henle’s loop; medullary thick ascending limb; cortical thick ascending limb; distal convoluted tubule; connecting tubule; cortical collecting duct; outer medullary collecting duct; inner medullary collecting duct.

RNAseq analysis of the undamaged right kidneys revealed *Kynu* as one of the top candidate genes significantly induced by HP and CR (Fig. 1B, Fig. S1A, B, http://shiny.cecad.uni-koeln.de:3838/IRaP) (18). Previously published single-cell (22), single-nucleus (23) and bulk-RNAseq (24) data as well as proteome analyses of microdissected kidneys (25) consistently all show predominant expression of *Kynu* in proximal tubules, a central site of damage susceptibility in AKI (Fig. 1C, Fig. S1C-F).

Immunohistochemical stainings of murine kidneys confirm this distinct localization of KYNU in the proximal tubuli (Fig. 1D). The same holds true for human kidney tissue based on human protein atlas data (proteinatlas.org) (26) (Fig. 1E). We then went on to confirm the preconditioning-mediated induction on the protein level by immunoblotting. Semiquantitative analyses reveal a significant increase of KYNU in response to CR but not to HP (Fig. 1F). Finally, in a mass spectrometry-based proteome dataset of kidneys after preconditioning, KYNU is detected in 8 out of 10 CR samples, but was never found after HP or in non-preconditioned controls (Fig. S1G, H).

### Generation of conventional KYNU^null^ mice

To further investigate the functional role of KYNU in HP/CR-mediated nephroprotection, KYNU-deficient (KYNU^null^) mice were generated by CRISPR/Cas9-mediated non-homologous end joining (NHEJ) (Fig. 2A). NHEJ resulted in a 2 bp-deletion leading to a frameshift and premature stop codon after 6 amino acids in exon 2 (Fig. 2B). KYNU-deficiency was successfully confirmed by immunoblotting (Fig. 2C), immunohistochemistry (Fig. 2D) and targeted liquid chromatography coupled to tandem mass spectrometry (LC-MS/MS) (Fig. S2).

**Fig. 2:**
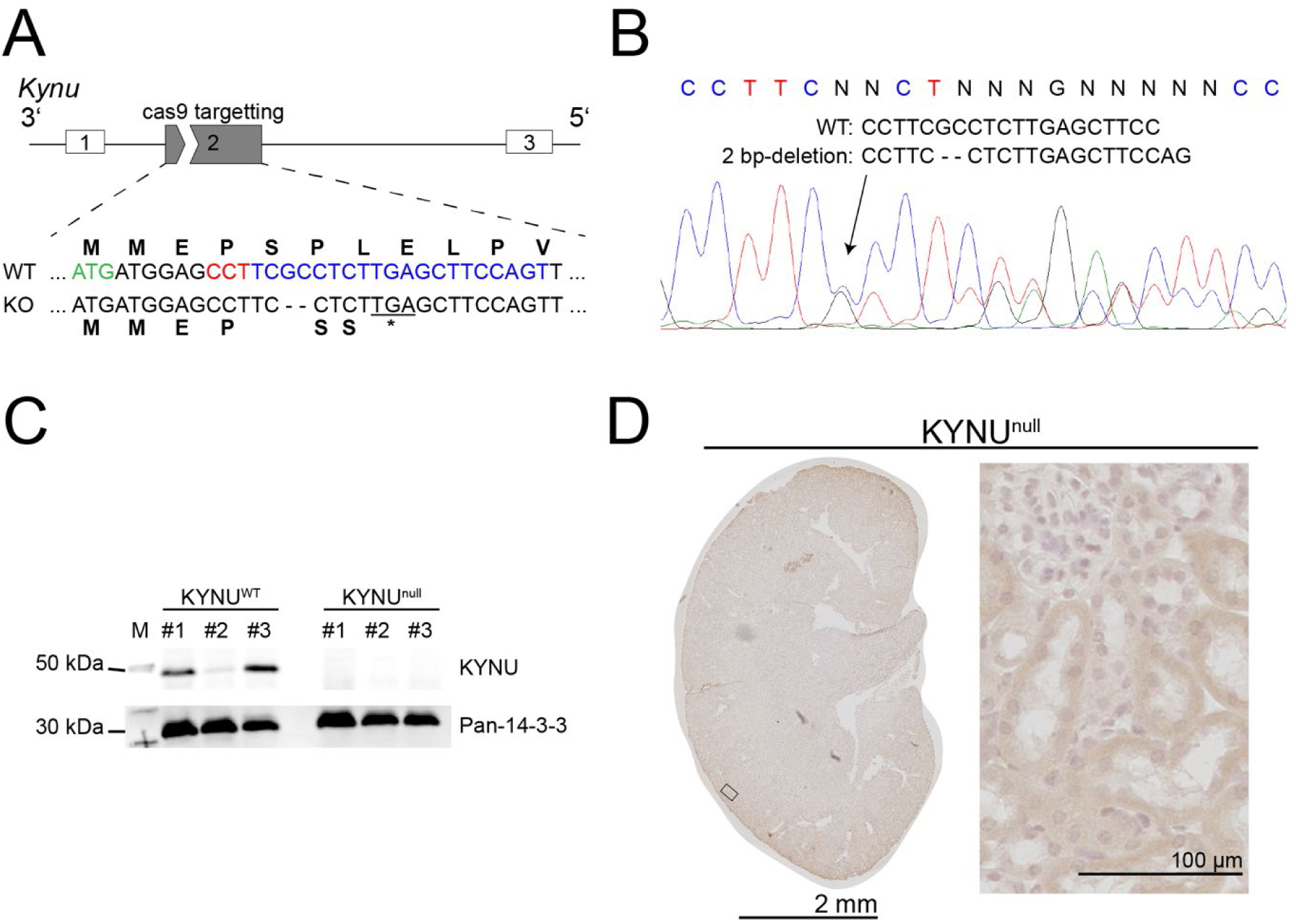
Generation of conventional KYNU^null^ mice. **A.** Schematic illustration of the strategy to generate KYNU^null^ mice by CRISPR/Cas9-mediated non-homologous end joining (NHEJ). **B.** Extraction of a Sanger sequencing displaying the 2 bp deletion (del) leading to a frameshift and premature stop codon after 6 amino acids in *Kynu*-exon 2. **C.** Immunoblot of murine kidneys showing the successful deficiency in KYNU^null^ mice compared to expression in WT mice (loading control: Pan-14-3-3). **D.** Histological analyses by staining with anti-KYNU in kidneys of KYNU^null^ mice confirming the deficiency compared to kidneys of WT (Fig. 1D). **Abb.:** WT: wildtype, KO: KYNU-deficiency; green: first methionine, red: PAM-side, blue: CRISPR-RNA sequence; bold capitals: aminoacids; KYNU^null^: KYNU-deficiency; KYNU^WT^: wildtype.

### KYNU^null^ mice are viable and do not show any abnormalities in general appearance or kidney function

Regarding lifespan (data not shown) and physical characteristics, i.e. external appearance and weight, (Fig. 3A, 3B), no differences are apparent in KYNU^null^ mice compared to wildtype (WT) littermates. Moreover, neither macroscopic nor microscopic examination of organ morphology of the kidney, liver, heart and brain reveals any alterations (Fig. S3). Detailed examination of kidney function shows no changes in plasma creatinine and blood urea nitrogen (BUN) levels as well as proteinuria in KYNU^null^ animals (Fig. 3C-F).

**Fig. 3:**
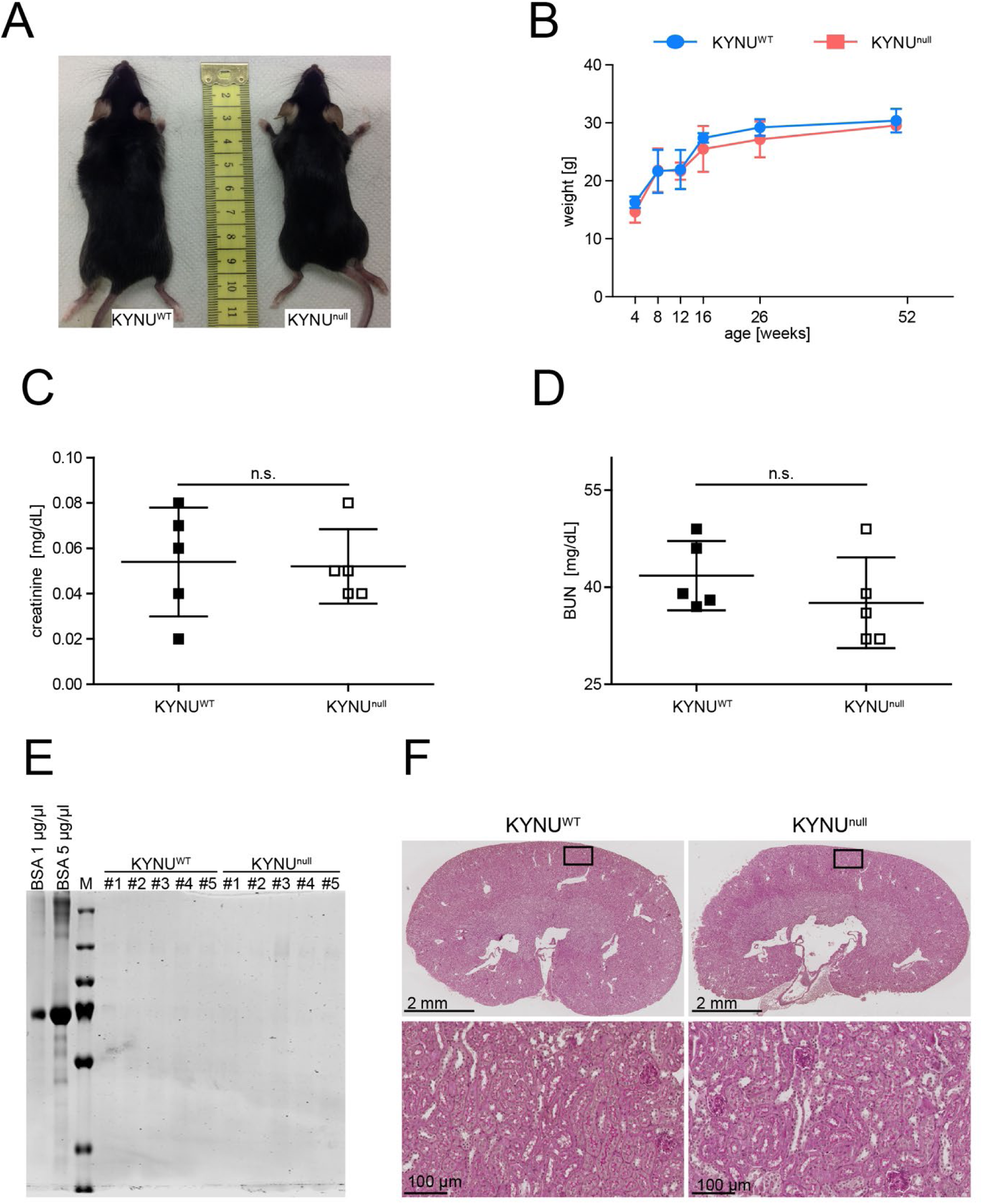
KYNUnull mice show a normal general phenotype and kidney function. A. Representative photograph of a wildtype (WT) mouse and a KYNUnull littermate showing no differences in general external appearance. B. Bodyweight of KYNUnull mice and wildtype littermates recorded for 1 year at the age of 4, 8, 12, 16, 26 and 52 weeks (n=5 per group). C. Plasma creatinine values of KYNUWT and KYNUnull mice; student’s t-test, n=5 per group. D. Plasma values of blood urea nitrogen (BUN) of KYNUWT mice and KYNUnull mice; student’s t-test, n=5 per group. E. Coomassie blue staining to screen for alterations of proteinuria in KYNUnull mice compared to WT mice (loading control: BSA) F. PAS staining of kidneys of a 14-week-old WT mouse and a 14-week-old KYNUnull mouse. Abb.: Bars indicate means ± standard deviation; BSA: bovine serum albumin; PAS: Periodic acid-Schiff reaction; WT: wildtype, n.s.: p-value≥0.05

### Loss of KYNU diminishes the protective potential of HP/CR

To examine whether KYNU is functionally involved in preconditioning-mediated nephroprotection, we performed preconditioning treatment followed by renal IRI in both KO animals and WT littermates (Fig. S4). 24 h after IRI, creatinine and blood urea nitrogen (BUN) of nonPC WT mice are significantly increased compared to sham-operated controls (sham WT, Fig. 4A, 4B). HP significantly reduces (p<0.01) and CR almost entirely prevents (p<0.0001) this marked increase of creatinine and BUN in WT animals (Fig. 4A, 4B). KYNU-deficiency does not change the outcome in non-preconditioned animals (nonPC KYNU^null^, sham KYNU^null^). However, protection by HP and CR is reduced in KYNU^null^ mice (Fig. 4A, 4B). Here, protection by HP does not reach statistical significance and KYNU-deficient CR mice show significantly higher levels of creatinine (p<0.05) and BUN (p<0.05) after IRI compared to corresponding WT littermates (Fig. 4A, B).

**Fig. 4:**
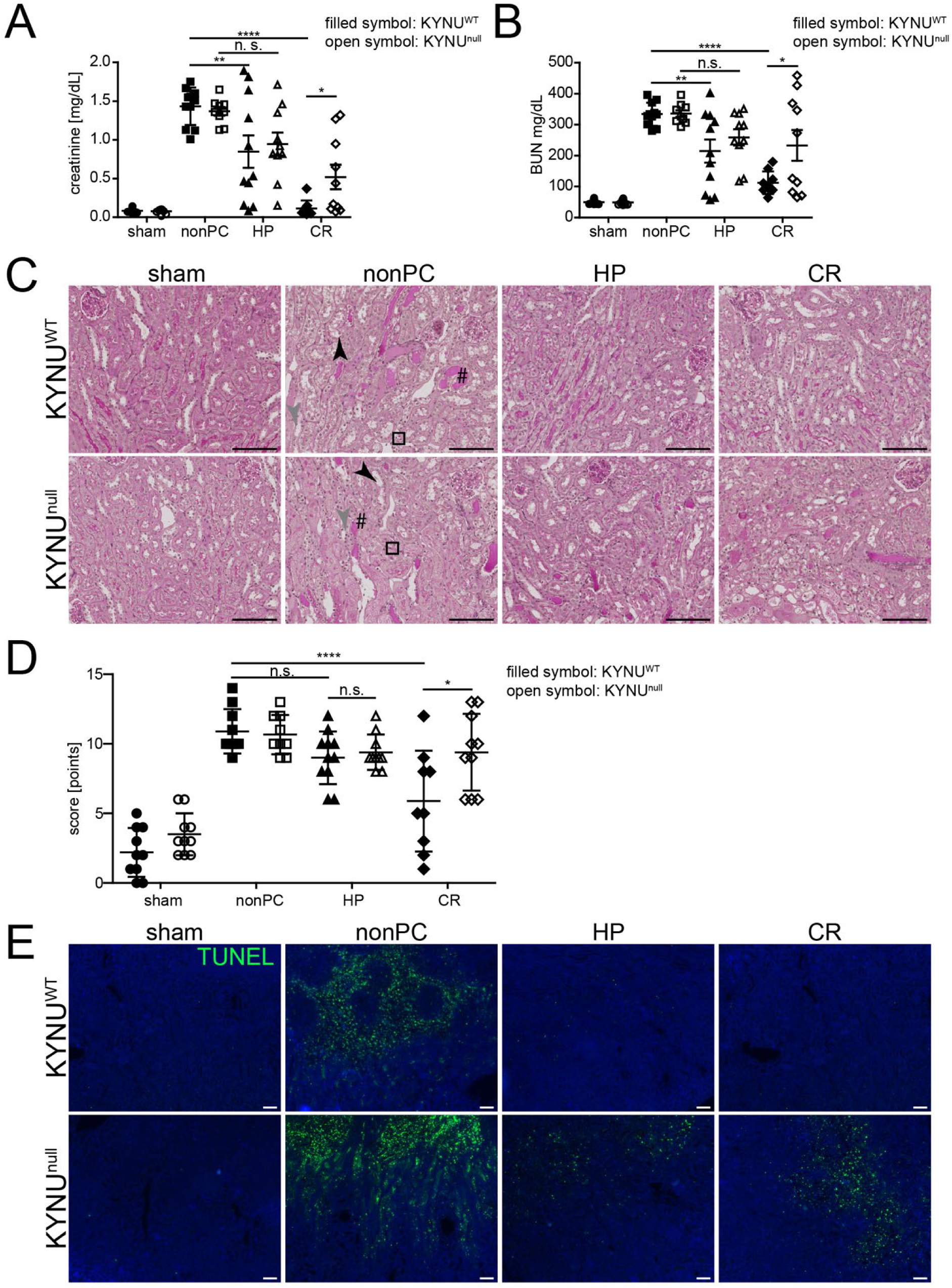
Loss of KYNU does not lead to kidney damage after IRI but diminishes the protective potential of preconditioning**. A.** Plasma creatinine values 24 hours after IRI. IRI induces a similar increase of plasma creatinine in KYNU^null^ and WT mice. KYNU-deficiency leads to a reduction of the protection by CR and shows a similar trend regarding HP. NonPC, HP and CR were analyzed using one-way ANOVA and Tukey’s posthoc test for multiple comparisons. For the comparison of HP-KYNU^WT^ and HP-KYNU^null^ and CR-KYNU^WT^ and CR-KYNU^null^ mice, students’ t-test was used. **B.** Plasma values of blood urea (BUN) nitrogen 24 hours after IRI recapitulate the findings for plasma creatinine. NonPC, HP and CR were analyzed using one-way ANOVA and Tukey’s posthoc test for multiple comparisons. For the comparison of HP-KYNU^WT^ and HP-KYNU^null^ and CR-KYNU^WT^ and CR-KYNU^null^ mice, students’ t-test was used. **C.** Periodic acid-Schiff reaction (PAS) of kidneys 24 hours after IRI. #: protein casts; square: pyknosis; black arrowhead: loss of brush border; gray arrowhead: epithelial flattening. **D.** Semiquantitative composite damage score; one-way ANOVA and Tukey posthoc test (n=10-12 per group) **E.** Analysis of cell death 24 hours after IRI using terminal deoxynucleotidyl transferase dUTP nick end labeling (TUNEL). HP and CR strongly protect from cell death whilst this potential is partly lost in KYNU^null^ mice. **Abb.:** Bars indicate means ± standard deviation; nonPC: non-preconditioned male mice; HP: hypoxic preconditioning; CR: caloric restriction; IRI: ischemia-reperfusion injury; KYNU^null^: KYNU-deficiency; KYNU^WT^: wildtype; sham: right nephrectomy followed by no-clamping of the left renal pedicle; scale bars indicate 100 µm, ****: p-value<0.0001; ***: p-value<0.001; **: p-value=0.001-0.01; *: p-value<0.05; n.s.: p-value≥0.05.

Periodic Acid-Schiff (PAS) staining confirms the involvement of KYNU in nephroprotection. Kidneys after IRI of nonPC WT and KYNU mice show increased pyknosis, stronger brush border loss and greater epithelial flattening compared to sham-operated mice (sham KYNU^WT^, sham KYNU^null^) (Fig. 4C). There is no difference in histological damage between KYNU^WT^ and KYNU^null^ animals in non-preconditioned animals. However, in WT mice, HP tends to reduce tissue damage and CR efficiently prevents histological signs of injury (HP/CR KYNU^WT^, Fig. 4C). After IRI the protective effect of CR is almost lost in KYNU^null^ mice (Fig. 4C). This was verified in a semi-quantitative analysis of the histological damage using a composite damage scoring system (Fig. 4D). Whilst protection by HP does not reach statistical significance, CR clearly prevents (p<0.001) this marked increase of the histological damage in KYNU^WT^ mice. However, the protective capacity of CR is lost in KYNU-deficient animals (Fig. 4D).

Terminal deoxynucleotidyl transferase dUTP nick end labeling (TUNEL) highlighting cell death support the histological findings (Fig. 4E). Whilst damaged kidneys of the nonPC group show strong signals in WT as well as KYNU^null^ mice (nonPC WT/KYNU^null^), only slightly positive signals comparable to sham (sham WT, sham KO) are detectable in preconditioned WT animals (HP WT, CR WT). Conversely, a marked TUNEL positive signal is observed in preconditioned KYNU^null^ mice (HP KO, CR KO) and is only slightly ameliorated compared to non-preconditioned animals.

### CR modulates the tryptophan metabolic pathway in a KYNU-dependent manner

To gain insights into the molecular mechanism underlying the impact of CR as well as KYNU on organ protection, we performed targeted metabolomics analyses of metabolites in the tryptophan metabolic pathway (pathway shown in Fig. 5A).

**Fig. 5:**
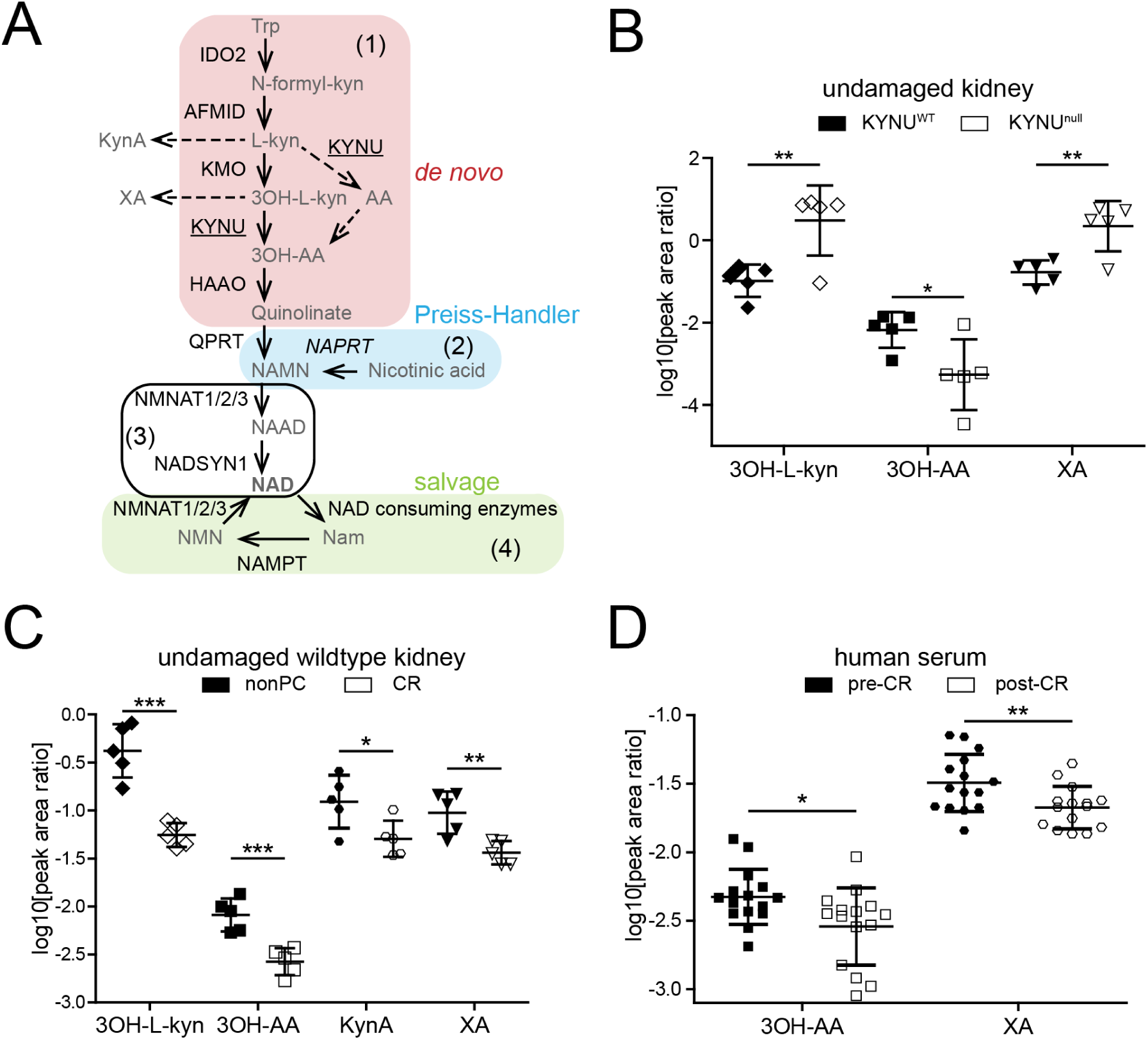
CR modulates the tryptophan metabolic pathway in a KYNU-dependent manner **A.** Illustration of the tryptophan metabolic pathway as one of the three NAD+ biosynthesis pathways. (1): *de novo*; (2): Preiss-Handler; (3): belonging to 1, 2 and 4; (4): salvage. **B.** Quantification of metabolites in the tryptophan metabolic pathway comparing undamaged kidneys of KYNU^WT^ and KYNU^null^ mice (unpaired student’s t-test, n=5 per group). Only metabolites that show a significant change are shown in this graph (for others see Suppl. Fig. S5). **C.** Quantification of metabolites involved in the tryptophan metabolic pathway comparing WT kidneys from the nonPC and CR groups (unpaired student’s t-test, n=5 per group). Only metabolites that show a significant change are given (for others see Suppl. Fig. S5). **D.** Quantification of metabolites involved in the tryptophan metabolic pathway comparing longitudinally acquired human serum samples pre- and post-CR (paired student’s t-test,n=15 per group). Only metabolites that show a significant change are given (for others see Suppl. Fig. S5). **Abb.:** (1): *de novo* branch of NAD+ biosynthesis; (2): Preiss-Handler branch of NAD+ biosynthesis; (3): belonging to 1, 2 and 4; (4): salvage branch of NAD+ biosynthesis; Trp: Tryptophan; IDO2: Indoleamine 2,3-dioxygenase 2; N-formyl-kyn: N-formyl-Kynurenine; AFMID: Arylformamidase; L-Kyn: L-Kynurenine; KynA: Kynurenic acid; KYNU: Kynureninase; AA; Anthranilic acid; KMO: Kynurenine 3-monooxygenase; 3OH-L-kyn: 3-Hydroxy-L-kynurenine; XA: Xanthurenic acid; 3OH-AA: 3-Hydroxy-anthralinic acid; HAAO: 3-Hydroxyanthranilate 3,4-dioxygenase; QPRT: Quinolinate Phosphoribosyltransferase; NAPRT: Nicotinate phosphoribosyltransferase; NAMN: Nicotinic acid mononucleotide; NMNAT 1/2/3: Nicotinamide mononucleotide adenylyltransferase 1/2/3; NAAD: nicotinic acid adenine dinucleotide; NADSYN1: NAD Synthetase 1; NAD: Nicotinamide adenine dinucleotide; NAM: nicotinamide; NAMPT: nicotinamide phosphoribosyltransferase; NMN: nicotinamide mononucleotide; KYNU^WT^: wildtype; KYNU^null^: KYNU-deficiency; nonPC: no preconditioning; CR: caloric restriction; bars indicate means ± standard deviation; ***: p-value<0.001; **: p-value: 0.001-0.01; *: p-value<0.05; n.s.: p-value ≥0.05.

Comparative analyses of undamaged, non*-*preconditioned kidneys of the two mouse lines reveal significant changes in metabolite abundance. While tryptophan (Trp) and L-kynurenine (L-kyn) levels are not affected by loss of KYNU (Fig. S5A), KO animals show a significant increase of 3-Hydroxy-L-kynurenine (3OH-L-kyn) (p<0.01) and a significant decrease of 3-Hydroxy-anthralinic acid (3OH-AA) (p<0.05) (Fig. 5B). This finding is in line with the function of KYNU, which is known to catalyze the reaction from 3OH-L-kyn to 3OH-AA (27). Correspondingly, in kidneys KYNU*-*deficiency leads to a significant increase of Xanthurenic acid (XA) – a known alternative metabolite of 3OH-L-kyn (p<0.01) and the formation of which does not depend on KYNU (Fig. 5B). Analyses of liver tissue reveal even stronger increases in the levels of 3OH-L-kyn (p<0.001) and XA (p<0.001) in KYNU^null^ mice (Fig. S5B). All findings can be recapitulated in measurements from blood plasma and a significant increase of Kynurenic acid (KynA) (p<0.05) is detected in blood plasma (Fig. S5C). In urine, 3OH-L-kyn and XA levels are also increased and 3OH-AA levels reduced (Fig. S5D). Finally, urinary Trp concentration is lower in KYNU^null^ animals, but urinary L-kyn levels remain unchanged by loss of KYNU (Fig. S5D).

Based on the induction of *Kynu* by HP and CR, opposite effects of both preconditioning strategies on tryptophan metabolites compared to KYNU-deficiency would be expected. In line with this hypothesis, CR does indeed induce significant reductions of 3OH-L-kyn (p<0.001), KynA (p<0.05) and XA (p<0.01) (Fig. 5C). Interestingly, 3OH-AA (p<0.01) is also reduced by CR (Fig. 5C). HP diminishes only 3OH-L-kyn (p<0.01) without significant effects on any other metabolite examined (Fig. S5E).

To evaluate whether the CR-induced metabolic effects in mice are conserved in humans, targeted metabolomics analyses were repeated in human samples. These specimens were derived from a cohort that had undergone CR in a previously published investigator-initiated pilot study at our center (28) (CR_KCH, NCT01534364). As in mice, CR significantly reduces levels of 3OH-AA (p<0.05) and XA (p<0.01) (Fig. 5D), while no effects on other tryptophan metabolites are observed (Fig. S5F).

### CR induces profound changes in transcript and protein levels of enzymes involved in NAD+ biosynthesis and maintains NAD+ levels after renal IRI

Considering the role of the tryptophan metabolic pathway in NAD+ biosynthesis as well as the growing evidence on the role of NAD+ in nephroprotection, we used our previously published RNAseq (18) and newly generated proteomics datasets to analyze renal transcript and protein levels of the key enzymes involved in the three NAD+ biosynthesis pathways as well as their modulation by IRI and preconditioning (Table S1). On the transcript level, CR leads to a significant modulation of genes involved in *de novo* NAD+ biosynthesis (tryptophan metabolic pathway), including the induction of *Kynu* described in Figure 1 (pre-IRI, log2FC: 2.16; adjusted p: 4.05*10-31). Transcript levels of Arylformamidase (*Afmid*) (log2FC: -0.33; adjusted p: 0.028) and 3-Hydroxyanthranilate 3,4-dioxygenase *(Haao)* (log2FC: -0.34; adjusted p: 8.5*10-05) are slightly, but significantly, reduced (Fig. 6A). In contrast, all changes in transcripts belonging to the Preiss-Handler or salvage branch of NAD+ biosynthesis by CR do not reach statistical significance. Further, in comparison to nonPC kidneys pre-IRI, HP does not significantly change the RNA expression of genes involved in NAD+ biosynthesis except *Kynu* (log2FC: 0.53; adjusted p: 2.7*10-4) (Fig. S6A).

**Fig. 6:**
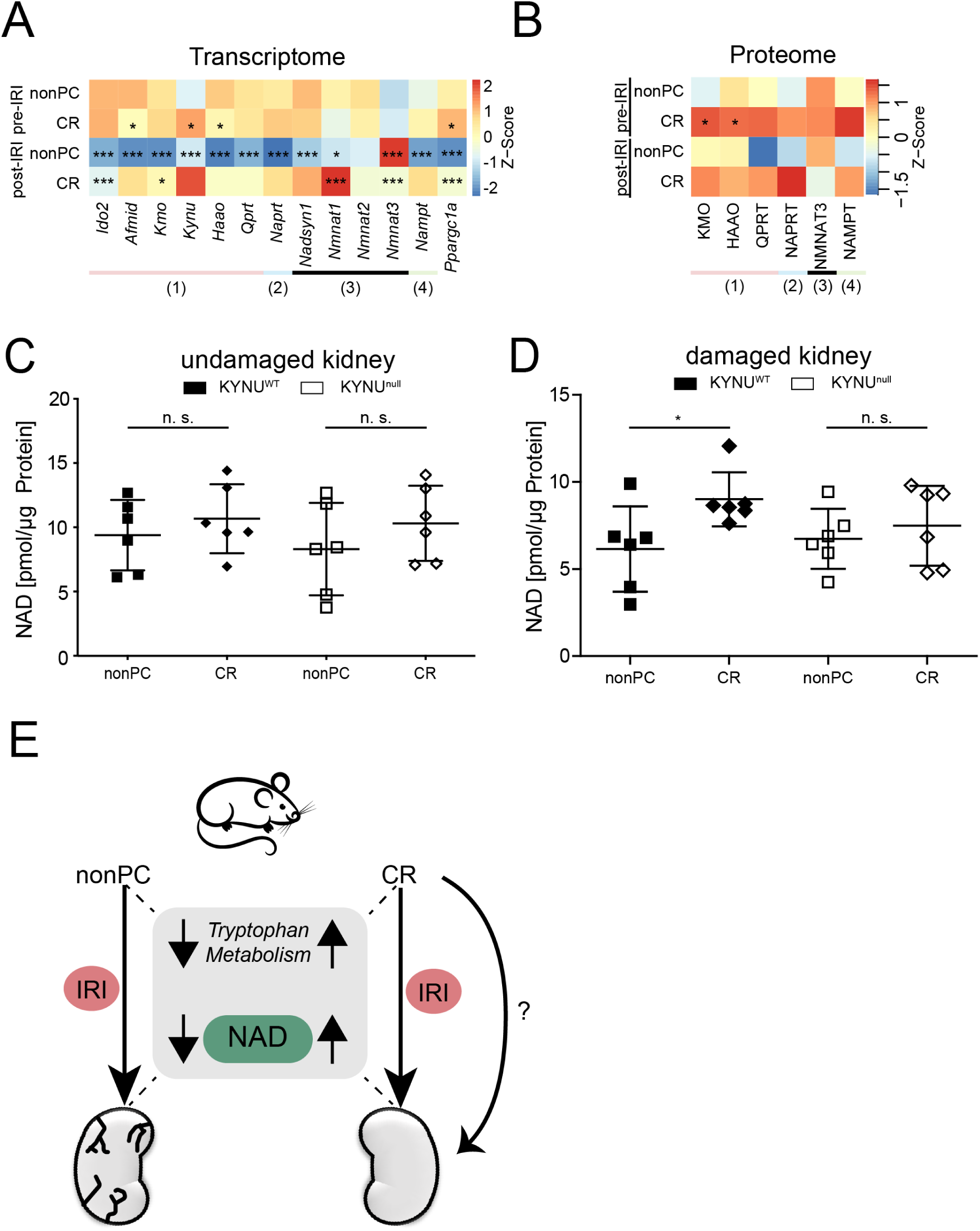
CR induces profound changes in transcript and protein levels of genes involved in NAD+ biosynthesis and maintains NAD+ levels after renal IRI **A.** Heatmap depicting mean based z-score transcript levels (RNAseq) of the genes involved in the NAD+ biosynthesis pre-IRI and at 24 h post-IRI in non-preconditioned (nonPC) mice or after CR; asterisks reflect significance of pairwise comparisons of each gene between treatment groups (n=5 per group). **B.** Heatmap depicting median based z-score protein expression levels (LC-MS/MS) of the enzymes involved in NAD+ biosynthesis (selection since not all proteins were detected by LC-MS/MS) at 0 h prior to (pre-IRI) and at 4 h after IRI (post-IRI) in non-preconditioned (nonPC) mice or after CR; asterisks reflect significance of pairwise comparisons of each gene between treatment groups (n=5 per group). **C.** NAD levels of right, undamaged kidneys of KYNU^WT^ and KYNU^null^ mice without previous preconditioning (nonPC) or after CR (student’s t-test, n=6 per group). **D.** NAD levels of left, damaged kidneys 24 h after IRI of KYNU^WT^ and KYNU*^n^*^ull^ mice without previous preconditioning (nonPC) or after CR (student’s t-test, n=6 per group). **E.** Model of the interaction between CR-mediated nephroprotection and *de novo* NAD+ biosynthesis (tryptophan metabolic pathway). The tryptophan metabolic pathway is activated by CR. In the context of IRI, this activation leads to a preservation of NAD levels and protects from AKI. Besides, CR might also exert some still unknown effects independent of the tryptophan metabolic pathway on the kidney (indicated by the interrogation mark). **Abb.:** (1): *de novo* branch of NAD+ biosynthesis; (2): Preiss-Handler branch of NAD+ biosynthesis; (3): belonging to 1, 2 and 4; (4): salvage branch of NAD+ biosynthesis; nonPC: non-preconditioned male mice; CR: caloric restriction;IRI: ischemia-reperfusion injuryNAD: total NAD (NAD+ & NADH); n. s.: p-value>0.05, *: adjusted p-value/q-value<0.05, *** adjusted p-value/q-value<0.0001

In non-preconditioned animals, IRI (24 h) significantly reduces the expression of nearly all the 12 genes involved in the three NAD+ biosynthesis pathways except Nicotinamide Nucleotide Adenylyltransferase 2 (Nmnat2) which is not significantly changed (adjusted p: 0.8) and Nmnat3, which is significantly induced (log2FC: 1.02; adjusted p: 3.1*10-14) by IRI. (Fig. 6A). Notably, CR prevents the IRI-induced changes of transcript levels for most of these genes and only Indoleamine 2,3-dioxygenase 2 *(Ido2)*, Kynurenine 3-monooxygenase (*Kmo)* and Nicotinamide Nucleotide Adenylyltransferase 1 (*Nmnat1*) are significantly regulated after IRI in CR-preconditioned animals. Whilst *Ido2* (log2FC: -1.28, adjusted p: 6.06*10-13) and *Kmo* (log2FC: - 0.3; adjusted p: 0.04) are significantly reduced, *Nmnat1* shows an opposite regulation by IRI in CR animals compared to the nonPC group (log2FC: 0.6; adjusted p: 4.9*10-6). Additionally, comparing transcript expression levels of CR kidneys to non-preconditioned (nonPC) kidneys post-IRI, *Nmnat3* shows a significantly lower abundance in CR than in non-preconditioned mice (log2FC: -0.71; adjusted p: 8.2 *10-5) (Fig. 6A). Post-IRI HP maintained transcript expression levels of Glutamine-dependent NAD+ synthetase (*Nadsyn1)*, *Nmnat1/2* as well as Nicotinamide phosphoribosyltransferase (*Nampt) compared to pre-IRI*. Similar to CR, *Nmnat3* (log2FC: 0.5; adjusted p: 2.0*10-3) is significantly induced by HP (Fig. S6A). RNA expression of Peroxisome proliferator activated receptor gamma coactivator 1 alpha *(Ppargc1α),* a key regulator of NAD+ biosynthesis, was slightly induced by CR (log2FC: 0.2*, adjusted p: 2.4*10^-4)* but without significant changes upon HP (log2FC: 0.017, adjusted p: 0.96). Post-IRI, *Ppargc1α* is reduced under all conditions compared to pre-IRI, whereas the expression in CR samples is significantly higher than in non-preconditioned samples (log2FC: 0.9, adjusted p: 4.48*10-11).

As expected, changes on the proteome level are much milder than the ones observed regarding transcript abundance. Out of the 12 enzymes analyzed, 6 could be identified and quantified by mass spectrometry (Fig. 6B). Effects are detected only in the *de novo* branch of NAD+ biosynthesis (Table S2). KYNU was measured in 3 out of 5 samples after CR pre-IRI but in none of the nonPC or HP samples, thereby underlining its CR-mediated induction. Pre-IRI CR induces slightly higher protein abundance for KMO (log2FC: 0.5, q-value: 0.018) and HAAO (log2FC: 0.53, q-value: 0.02), whilst the protein expression levels of Quinolinate Phosphoribosyltransferase (QPRT, log2FC: 0.22, q-value: 0.1), Nicotinate phosphoribosyltransferase (NAPRT, log2FC: 0.17; q-value: 0.18) and NMNAT3 (log2FC: 0.027; q-value 0.69) are not changed. IRI leads to a decrease of nearly all studied enzymes in CR samples (4 h after reperfusion) except KYNU (detectable in 5/5 CR-samples post IRI) and NAPRT (log2FAC: 0.07; q-value: 0.93), without a significant change of abundance. However, the IRI-mediated decrease in protein levels is again much attenuated by CR. In comparison to non-preconditioned samples, all targets except NMNAT3 show higher abundances after CR, indicating that the tryptophan metabolic pathway is preserved (Fig. 6B). Finally, as for the transcript levels, the effect of HP on protein abundance are much less pronounced than the effect seen for CR (Fig. S6B).

Considering the impact on key enzymes of NAD+ biosynthesis as well as the contribution of KYNU to the protective effect of CR regarding renal IRI, we went on to quantify NAD in both WT and KO animals. NAD levels of the undamaged right and damaged left kidneys of the same animals that were used for the IRI experiments (Fig. 4) were determined enzymatically using a colorimetric assay. Whilst NAD levels were not changed by CR in undamaged kidneys (Fig. 6C), kidneys after IRI showed significantly higher NAD levels in CR-treated compared to nonPC WT animals (Fig. 6D). Intriguingly, this effect of CR was entirely lost in KYNU*^null^* mice (Fig. 6D). Taken together, CR modulates *de novo* NAD biosynthesis (tryptophan metabolism pathway) leading to the maintenance of NAD levels after IRI. This CR-associated modulation of the tryptophan metabolic pathway is a mediator of the protective effects of preconditioning (Fig. 6E).

## Discussion

Preconditioning-mediated organ protection by transient HP and short-term CR has been shown to be extremely efficient in the context of IRI especially in rodents (14, 18). Nevertheless, the underlying molecular mechanisms modulating organ protection currently remain elusive. Recently, a transcriptome analysis comparing gene expression signatures identified *Kynu* as a potential common target of preconditioning (18).

*Kynu* encodes Kynureninase (KYNU), a key enzyme involved in the tryptophan metabolic pathway and *de novo* NAD^+^ biosynthesis. KYNU catalyzes the cleavage of L-kynurenine (L-kyn) and 3OH-L-kyn into anthranilic acid (AA) and 3-Hydroxyanthranilic acid (3-OH-AA), respectively (27, 29). While KYNU is expressed in almost all mammalian organs (30), the tryptophan metabolic pathway has been shown to be highest in the brain, liver and kidney (31, 32). KYNU has also been shown to be involved in inflammatory, tumorous, cardiovascular as well as chronic kidney diseases (27, 33–36). As a central enzyme in the tryptophan metabolic pathway, KYNU contributes to *de novo* NAD+ biosynthesis. NAD+ is a crucial and essential cofactor orchestrating redox reactions (31) as well as a substrate for several enzymes contributing to cellular stress resistance such as sirtuins (21). Consequently, it is not surprising that studies have linked alterations of the *de novo* NAD+ biosynthesis pathway to the modulation of lifespan (37) and organ protection (32, 38). However, to our knowledge, no study has as yet elucidated the role of KYNU in the context of preconditioning and ischemia-reperfusion injury or explored the underlying molecular mechanisms through which preconditioning mediates organ protection.

Based on the induction of *Kynu* observed upon both HP and CR, KYNU^null^ mice were generated using CRISPR/Cas9-mediated NHEJ. As previously described in mice with a *Kynu*-null allele of exon 3 (39), KYNU^null^ mice kept on standard chow did not show any abnormalities regarding viability, fertility or lifespan.

To date, published data had remained partly controversial regarding the role of *de novo* NAD+ biosynthesis in organ protection (32, 37, 40, 41). However, Parikh and coworkers elegantly showed that enzymes involved in the *de novo* NAD+ biosynthesis were reduced in AKI in rodents (40). Interestingly, they observed reduced levels of quinolinate phosphoribosyltransferase (QPRT), necessary for nicotinate D-ribonuleotide synthesis from quinolate (QA), and consecutively, elevated levels of urinary QA in patients at risk for AKI. These findings indicate an important role of the tryptophan metabolic pathwayin human AKI. Moreover, PGC1α, the downstream co-activator of *de novo* NAD+ biosynthesis, plays a central role in the development of AKI (21, 38, 40). Nevertheless, non*-*preconditioned KYNU*^null^* mice did not show an altered outcome after renal IRI in our study – a finding that is in line with previously published data (37). Consequently, the *de novo* NAD+ biosynthesis pathway does not appear to contribute to protection from renal IRI under standard conditions as suggested by previous studies demonstrating that most NAD+ is generated by the salvage pathway (42, 43). However, KYNU-deficiency indeed partly abrogated the protective potential of CR. Consequently, *de novo* NAD+ biosynthesis through the tryptophan metabolic pathway is employed by preconditioning to protect from IRI.

Several intermediate metabolites of the kynurenine pathway, e.g. 3OH-L-kyn, 3OH-AA, XA and KynA, had previously been discussed as modulators of organ damage through their immunomodulatory, transcription regulating and antioxidant properties (31, 32, 44–46). In our study, KYNU-deficiency resulted in increased levels of 3OH-L-kyn as well as XA in tissue (kidney and liver), blood plasma and urine, whereas KynA was only increased in blood plasma and urine. Furthermore, KYNU-deficiency resulted in a reduction of 3OH-AA in the kidney as well as in urine. Since KYNU-deficiency did not alter kidney function after IRI, but rather increased 3OH-L-Kyn and XA at a basal level in non*-*preconditioned mice, this points towards a negligible role of these intermediates in kidney injury. The role of KynA may be more complex. Elevated KynA levels resulting from inhibition of α–ketoglutarate (aKG)-dependent dioxygenase 1 (Egln1) and an altered tryptophan metabolic pathway, i.e. reduced abundance of Trp and L-Kyn, is known to effectively protect from myocardial injury (44). Hypothetically, this protective effect could be explained by the tryptophan metabolic pathway exploiting increased KynA levels for *de novo* generation of NAD+. However, to our knowledge, metabolization of KynA via the tryptophan metabolic pathway in this direction has not as yet been shown. Moreover, this hypothesis does not directly explain the kidney protection achieved by Kynurenine 3-monooxygenase (KMO)-deficiency resulting in elevated KynA plasma levels (32). Since L-Kyn can be metabolized to 3OH-AA in a KMO-independent manner, KMO-deficiency does not completely interrupt the tryptophan metabolic pathway (Fig. 5A, (32). Therefore, it seems possible that KMO-deficiency leads to an induction of the tryptophan metabolic pathway as indicated by reduced 3OH-AA levels in *KMO^null^* mice without IRI in comparison to the stronger increase of 3OH-AA observed in *KMO*^null^ mice after IRI (32). In our study, KynA was elevated in blood plasma and urine in KYNU^null^ mice, but the increased abundance of KynA in KYNU^null^ mice was not sufficient to protect from IRI. The reason for this finding is currently not clear. However, it may be a consequence of a lack of increase in kidney tissue or due to the KYNU-deficiency hampering a transfer of KynA into the *de novo* NAD+ biosynthesis pathway.

CR strongly affected tryptophan metabolite levels, whereas reduced 3OH-L-kyn was the only effect detected after HP. This discrepancy is likely to be a consequence of the much less potent impact of HP on *Kynu* expression which may also be one of the reasons for the more profound protection evoked by CR. The fact that preservation of NAD+ after IRI evoked by CR was abrogated in KYNU^null^ mice shows that the *de novo* NAD+ biosynthesis pathway is involved in CR-mediated organ protection.

Gene expression analyses of key enzymes in *de novo* NAD+ biosynthesis showed that IRI was associated with the downregulation of these genes (e. g. *IDO2*, *Afmid*, *Kmo*, and *Kynu*) in non-preconditioned mice. Intriguingly, CR did not only maintain the levels of these enzymes, but IRI even induced *Kynu* and *Nmnat1*. *Nmnat1* belongs to the family of Nicotinamide Mononucleotide Adenylyl Transferases (NMNATs) consisting of NMNAT1-3. These enzymes show distinct subcellular localizations, e.g. NMNAT1 in the nucleus, NMNAT2 in the cytoplasma and the cytoplasm and NMNAT3 in the mitochondria (47–49). All three NMNATs (NMNAT1-3) are part of the salvage and *de novo* branches of NAD+ biosynthesis (31). Interestingly, *Nmnat2* and *3* were the only transcripts of enzymes for which the levels increased after IRI in non-preconditioned animals. In contrast, *Nmnat1* was only induced in CR-treated mice pointing to specific responses of subcellular compartments in CR-treated mice. *Ppargc1α*, a downstream mediator of *de novo* NAD+ synthesis and a transcriptional co-activator of mitochondrial biogenesis interacting with the nuclear peroxisome proliferator-activated receptor gamma (PPARγ) (38, 40), was increased after CR pre-IRI and its decrease post-IRI was reduced by CR. Thus, in calorically restricted mice, the induction of nuclear *Nmnat1* might result from an induction of *de novo* NAD+ synthesis, whilst in non-preconditioned kidneys, mitochondrial *Nmnat3* is induced in an attempt to preserve NAD+ by the salvage pathway. Taken together, CR has profound effects on *de novo* NAD+ biosynthesis and the resulting maintenance of NAD+ levels after IRI is an important contribution to preconditioning-mediated nephroprotection. It is therefore very likely that this mechanism is also involved in the effects of CR in other organs. However, considering that loss of KYNU does not completely abrogate the nephroprotection after CR, additional pathways or mediators could be involved in this preconditioning strategy. Importantly, identification and combination of all of these players will be important in future clinical translational studies.

Since translation of interventions that mediate organ protection to the human setting is highly desirable, a better fundamental understanding of the impact of preconditioning on metabolic pathways in humans is a prerequisite before translating to a clinical setting. Importantly, in this study, blood serum analysis of patients showed that the CR-induced signature of the tryptophan metabolic pathway is indeed conserved in humans - a finding that can be exploited in future clinical trials targeting NAD+ synthesis

In conclusion, this proof-of-principle study strongly suggests that CR employs induction of the *de novo* NAD+ pathway to protect from renal IRI and that KYNU appears to be one of the central targets of different preconditioning strategies. These findings coupled with the conservation of the metabolic impact of CR on the tryptophan metabolic pathway in humans makes targeted modulation of the *de novo* NAD+ pathway an extremely interesting topic for future studies in humans.

## Methods

### Animal procedures

Initially, mice were provided by Charles River and then further bred in the CECAD *in vivo* Research Facility, University Hospital of Cologne. From the beginning, the mice were kept under a daily 12 hour light and 12 hour dark rhythm with similar specific pathogen-free (SPF) conditions in group cages with up to 5 mice at the same time. The mice received ad libitum access to drinking water as well as ad libitum access to food provided by sniff (Soest, Germany) with the exception of caloric-restricted mice.

### Generation of guide RNAs

A commercial KIT (Thermofisher, Waltham, Massachusetts, USA, A29377) was used for guide RNA (gRNA) generation. As previously described (50), crRNA (Alt-R^TM^ crRNA, IDT, Coralville, Iowa, USA) and tracrRNA were resuspended to 100 μM in nuclease-free and sterile T_10_E_0.1_ buffer (10 mM Tris-HCl, 0.1 mM EDTA, embryo-tested water, Sigma, W1503, St. Louis, Missouri, USA) prepared as described (51). By subjection of 1:1 equimolar ratios of crRNA and tracrRNA to 95°C for 5 min followed by reduction of 5°C/min, crRNA:tracrRNA complexes (50 μM) were generated. T7 RNA polymerase mediated *in vitro* transcription (NEB, Frankfurt, Germany, E2040S) was used for sgRNA-generation from the pSpCas9(BB)-2A-GFP (PX458) plasmid (Addgene, Watertown, Massachusetts, USA, #48138) from F. Zhang (McGovern Institute for Brain Research, Department of Brain and Cognitive Sciences, Department of Biological Engineering, Massachusetts Institute of Technology, Cambridge, MA 02139, USA), column purified (Qiagen, Hilden, Germany, 217004) and the gRNA was stored at -80°C. The crRNA sequences are listed is the paragraph concerning oligonucleotides used in this study below.

### Generation of KYNU^null^ mice

The generation of KYNU^null^ mice was carried out in our in-house Transgenic Core Unit (CECAD)) using C57Bl6N mice from Janvier LABS (Le Genest-Saint-Isle, France). A detailed protocol has been published previously (50). Female mice were stimulated with gonadotropin for 3 days and afterwards mated with male mice. Following the design of a gRNA, further procedures were performed by the CECAD *in vivo* Research Facility, Cologne. There, the fertilized ovum was removed and the CRISPR/Cas9-reaction mix (crRNA, generic tracrRNA, cas9-mRNA, see below for oligonucleotide sequences) was injected. Subsequent to the microinjection in the pronucleus, the reimplantation in female mice was performed (52).

### Confirmation of KYNU-deficiency

Occurring mutations within the first three generations of newborn mice were sequenced by the Cologne Center for Genomics (CCG). An ensuing proof of KYNU-deficiency was performed by immunoblot and immunohistochemistry. For this, WT and KO mice were sacrificed and kidneys were removed and snap frozen or fixed in 4% formaldehyde for further analyses. An anti-KYNU antibody (MAB7389, R&D Systems, Minneapolis, Minnesota, USA) was used for immunoblot analyses according to the manufactures’ protocol in a dilution of 1:1000. As a loading control, we used anti-pan-14-4-4 antibody (Sc-629, Santa Cruz Biotechnology, Dallas, Texas, USA) in a dilution of 1:1000 according to the manufacturer’s protocol. Immunohistochemical confirmation of KO was carried out by staining anti-KYNU antibody (MAB7389, R&D Systems, Minneapolis, Minnesota, USA) in a dilution of 1:50 according to the manufacturer’s protocol. Images were acquired in a 20x magnification using a slidescanner SCN4000 (Leica Microsystems, Jena, Germany).

For further determinations of the desired genotype, we designed primers to detect the mutation (IDT DNA technologies, Coralville, Iowa, USA; sequences listed below) and used tissue of ear marking for the following PCR analyses.

### Oligonucleotides used in this study

*For gRNA generation:*

m*Kynu* crRNA forward primer: **5’- TAATACGACTCACTATAG** ACTGGAAGCTCAAGAG**- 3’**

m*Kynu* crRNA reverse primer: **5’- TTCTAGCTCTAAAAC** TCGCCTCTTGAGCTTCCAG**- 3’**

*For genotyping:*

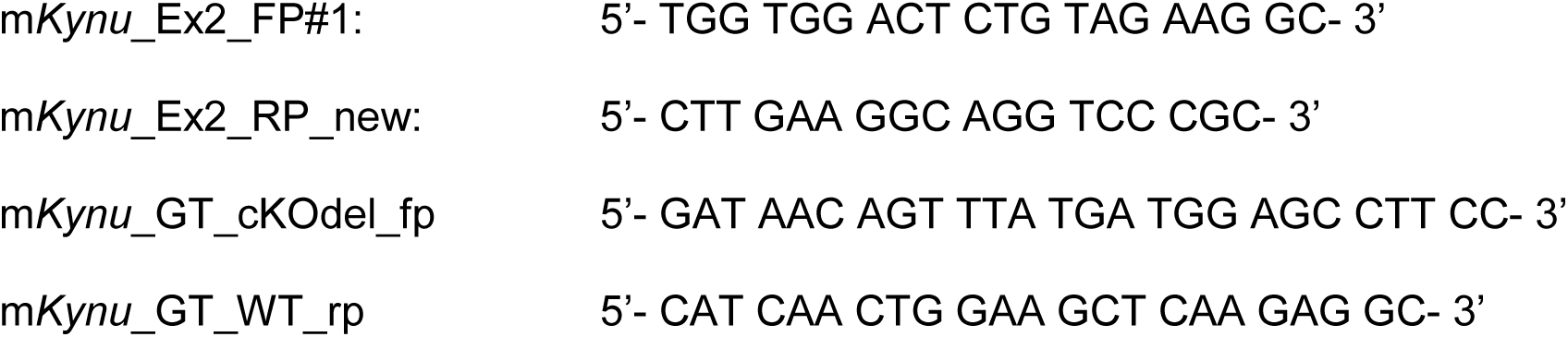

### Basal phenotyping

The experimental animals were weighed on a weekly basis. At the age of 12 weeks, a collection of urine was performed for further evaluation of the kidney function with the aid of coomassie blue staining. Beyond that, following an anesthesia by ketamin (Zoetis, Berlin, Germany) / xylazine (Bayer AG, Leverkusen, Germany) blood was taken in heparin-coated 1 ml syringes by cardiac puncture of the right ventricle for measurements of plasma urea and creatinine. In this context, both kidneys were removed and fixed in 4% formaldehyde or snap frozen in liquid nitrogen for further phenotyping.

### Preconditioning

For the hypoxic preconditioning, mice were kept in a hypoxia chamber with a reduction of oxygen to 8.3% for 2 h, 4 h and 8 h on 3 consecutive days prior to surgery based on the description of Bernaudin *et al.* (53).

For the caloric preconditioning, mice obtained 66% of the usual daily intake (3 g per day per mouse) 28 days before surgery as previously described (19, 54).

### Ischemia-Reperfusion Injury (IRI)

For the renal IRI surgery, we used 11- to 13-week-old male C57Bl6 WT mice and KYNU^null^ littermates (n=12 per group). Surgery was carried out on a temperature-controlled heating pad (Havard apparatus, Holliston, Massachusetts, USA) to keep the body temperature on one level. Following an anesthesia by ketamin (Zoetis, Berlin, Germany) / xylazine (Bayer, Leverkusen, Germany), mice received a midline abdominal incision as previously described (54). The vessels of the right kidney were ligated with sutures, and then the right kidney was removed and cut in two equal sections. One section was fixed in 4% formaldehyde and the other section was snap frozen in liquid nitrogen. After that the left renal artery and vein underwent unilateral clamping for 40 minutes by using an atraumatic microvascular clamp, followed by a subsequent removal of the clamp for reperfusion. The success of the reperfusion was controlled visually and afterwards the abdominal incision was closed in two layers by silk suture in the body wall and clips in the skin. The same procedure was performed on sham animals except for missing ischemic damage to the left kidney. Following the surgery process, all mice received normal saline containing 0.2 mg Buprenorphin subcutaneously and access to drinking water supplemented with Buprenorphin. Finally, the mice were placed into single cages to prevent mutilation of the wound by littermates. Following another anesthesia 24 h (4 h for proteomic analyses) after reperfusion, blood was taken in heparin-coated 1 ml syringes by cardiac puncture of the right ventricle. Afterwards, the blood was centrifuged at 4000 g for 10 minutes at 4°C. As done to the right kidneys before, the left kidneys were divided into two equal pieces. Again, one part was fixed in 4% formaldehyde and subsequently embedded in paraffin and the other part was snap frozen in liquid nitrogen for further analyses.

### Functional data and statistics

Accurate measuring of plasma urea and creatinine was performed by the central laboratory of the University Hospital of Cologne using a Cobas C 702 (Roche Diagnostics, Germany). Calculation of sample size was performed by P. S. on the basis on previous creatinine levels (effect size: 1.2; alpha error (two-sided): 0.05, power: 0.8). GraphPad Prism (GraphPad Software, San Diego, California, USA) was used for statistical analysis. For comparisons of more than two groups, one-way ANOVA followed by Tukey posthoc tests. For comparisons of two unpaired groups, two-tailed Student’s t-tests was used. In case of longitudinally acquired human samples, a paired two-tailed Student’s test was used. All functional data are represented as means ± standard deviations (SD) and two-tailed p-values<0.05 were considered significant. For transcriptomic analyses, adjusted p-values (padj)<0.05 were considered significant.

### Histopathology

After removal and fixation in 4% formaldehyde, all kidney organ tissue (kidney, liver, heart and brain) was embedded in paraffin and sliced into 2 µm sections. Using standard protocols for hematoxylin-eosin (HE), Periodic acid-Schiff (PAS) reaction and cresyl violet (CV) stainings, the slices were prepared for morphological assessments.

We involved an experienced nephropathologist to analyze the tubular damage of each kidney in a blinded fashion. He used a composite score as previously described (19, 55), which included the percent of tubules that presented increased pyknosis, stronger brush border loss and greater epithelial flattening and vacuolization. The mean for each treatment group was graphically represented. Moreover, terminal deoxynucleotidyl transferase dUTP nick end labeling (TUNEL) staining was carried out by using the DeadEnd Fluorometric TUNEL System (Promega, Fitchburg, Wisconsin, USA) according to the manufacturer’s protocol.

### Image acquisition

Images of PAS, HE, Kresylviolett and immunohistochemistry stainings were acquired in a 20x magnification using a slidescanner SCN4000 (Leica Microsystems, Jena, Germany). Images of immunofluorescence (TUNEL) were acquired by an AxioCamMR 702 camera on a Zeiss AxioObservermicroscope employing a Plan Apochromat 20x/0.8 objective (all from Carl Zeiss MicroImaging GmbH, Jena, Germany) supplied with ZEN software (version Zen 2.6, Carl Zeiss AG, Jena, Germany).

### Sample preparation for proteomic analysis using liquid chromatography with tandem mass spectrometry

Samples were prepared as previously described (56). In brief, kidney samples were homogenized and sonicated in urea buffer (8 M), ammonium bicarbonate (100 mM) and protease inhibitor (Roche Diagnostics GmbH, Penzberg and Mannheim, Deutschland). Each sample was reduced with dithiothreitol and alkylated using iodoacetamide afterwards. Overnight proteins were digested using trypsin (1:100 w/w ratio) and then cleaning of peptides was performed by use of StageTips (57). Afterwards, nano-liquid chromatography with tandem mass spectrometry was performed on a Q Exactive Plus hybrid quadrupole-Orbitrap mass spectrometer (Thermo Fisher Scientific Inc., Waltham, MA) coupled to a nano-liquid chromatography device (Proxeon Odense, Syddanmark, Denmark) as previously described (58). For the separation of peptides, a 4-h gradient was used.

### Sample preparation for proteomics analysis using targeted mass spectrometry (PRM Assay)

Preparation of the samples for targeted mass spectrometry was performed as previously described (56). Livers of KYNU^WT^ and KYNU^null^ mice were homogenized and sonicated in a buffer containing 8 M urea, 100 mM ammonium bicarbonate and protease inhibitor (Roche, Penzberg, Germany). Afterwards, the samples were reduced using DTT and alkylated with IAA. Subsequently, proteins were digested with trypsin at a 1:100 w/w ratio at room temperature overnight. After that, peptides were cleaned up with stagetips (57). A peptide library generated from previously described mouse podocyte (59) and kidney cortex data (19) was generated using Skyline (60). Peptides were prioritized based on dot product and overall peak shape and the PRM Assay for protein identification (Peptide R.VAPVPLYNSFHDVYK.F++, RT = 37.2 min; loading control: ACT b, Peptide R.HQGVMVGMGQK.D++ RT = 8.9-13.4 min) was performed on a QExactive plus (Thermo Fisher Scientific Inc., Waltham, Massachusetts, USA) Orbitrap machine coupled to a nLC device (Proxeon AS, 5000 Odense, Denmark) as described before (61) using a 1-h gradient for separation of the peptides.

### Sample processing protocol for proteomic analyses

Samples were prepared as previously described (56). In brief, kidney samples were homogenized and sonicated in urea buffer (8M), ammonium bicarbonate (100 mM) and protease inhibitor (Roche Diagnostics GmbH, Penzberg and Mannheim, Deutschland). Each sample was reduced with dithiothreitol and alkylated using iodoacetamide afterwards. Proteins were digested overnight using trypsin (1:100 w/w ratio). Cleaning of peptides was performed by use of StageTips (57). Afterwards, nano-liquid chromatography with tandem mass spectrometry was performed on a Q Exactive Plus hybrid quadrupole-Orbitrap mass spectrometer (Thermo Fisher Scientific Inc., Waltham, Massachusetts, USA) coupled to a nano-liquid chromatography device (Proxeon Odense, Syddanmark, Denmark) as previously described (58). For the separation of peptides, a 4-h gradient was used.

### Data processing protocol for proteomic analyses

Raw data were searched against the Swissprot canonical mouse reference database (download from uniprot.org 03/2021). Data was analyzed using MaxQuant version 1.6.17.0 with the parameters uploaded with this submission. Data was further analyszed using Perseus 1.6.15.0 and 1.5.5.3. In brief, contaminants and reverse hits, as well as proteins only identified by site were removed. Protein label-free quantification values (MaxLFQ values) were log2 transformed and the data was filtered for the presence of at least 8 valid values per treatment group (group size: 10 samples), followed by median subtraction normalization.

### Quantification of metabolites of the tryptophan metabolic pathway

Selected metabolites of the L-tryptophan-kynurenine pathway were quantified by Liquid Chromatography coupled to Electrospray Ionization Tandem Mass Spectrometry (LC-ESI-MS/MS) using a procedure previously described (62) with several modifications.

Mouse kidney and liver samples were lyophilized overnight. After addition of ice-cold methanol/water 4:1 (v/v) / 0.2% formic acid (40 µl/mg dry weight), the tissue was homogenized using the Precellys 24 Homogenisator at 6,500 rpm for 30 sec and directly put on ice again. From this, 100 µl homogenate were added to 400 µl of ice-cold methanol/water 4:1 (v/v) / 0.2% formic acid and isotope-labeled internal standards (10 µl 2 µM ^13^C_3_,^15^N-3-hydroxykynurenine (Sigma-Aldrich) and 10 µl 2 µM D5-kynurenic acid (Eurisotop)). From human serum and mouse plasma and urine, 50 µl were added to 200 µl methanol, 0.2 % formic acid and the internal standards mentioned above. For urine samples, an internal standard mixture consisting of 5 µl 10 µM ^13^C_3_,^15^N-3-hydroxykynurenine, 5 µl 20 µM D5-kynurenic acid and 5 µl 10 µM D3-creatinine was used. After thorough mixing, the samples were incubated at -20°C for 30 min and centrifuged (16,100 RCF, 5 min, 4°C). The supernatant was dried under a stream of nitrogen and the residue was resolved in 100 µl 1% acetonitrile / 0.2% formic acid. After mixing and centrifugation, 90 µl supernatant were transferred to autoinjector vials and immediately measured.

LC-MS/MS analysis was performed by injection of 5 µl sample onto an Acquity UPLC BEH Shield RP18 column (100 mm × 2.1 mm ID, 1.7 µm particle size, 130 Å pore size, Waters) and detection using a QTRAP 6500 triple quadrupole/linear ion trap mass spectrometer (SCIEX). The LC (Nexera X2 UHPLC System, Shimadzu) was operated at 40°C and at a flow rate of 0.45 ml/min with a mobile phase of 1% formic acid in water (solvent A) and 0.2% formic acid in acetonitrile (solvent B). Metabolites were eluted with the following gradient: initial, 10% B; 2.60 min, 99% B; 3.19 min, 99% B; 3.20 min, 10% B; and 5.50 min, 10% B. Metabolites were monitored in the positive ion mode with their specific Multiple Reaction Monitoring (MRM) transitions (Table S3). The instrument settings for nebulizer gas (Gas 1), turbogas (Gas 2), curtain gas and collision gas were 50 V, 60 V, 40 V and medium, respectively. The Turbo V ESI source temperature was 450°C and the ionspray voltage was 5.5 kV. The values for declustering potential (DP), entrance potential (EP), collision energy (CE) and cell exit potential (CXP) of the different MRM transitions are listed in Table S3.

The quantifier peaks of endogenous metabolites and internal standards were integrated using the MultiQuant 3.0.2 software (SCIEX). The peak areas of the endogenous metabolites were normalized to those of the internal standards. The peak area of D5-kynurenic acid was used to normalize the peak areas of endogenous kynurenic acid, 3-hydroxyanthranilic acid and xanthurenic acid. ^13^C_3_,^15^N-3-hydroxykynurenine was used to normalize 3-hydroxykynurenine, tryptophan and kynurenine. For urine samples, these peak area ratios were additionally normalized to the peak area ratio of endogenous creatinine to D3-creatinine.

### NAD+/NADH quantification

For quantification of total NAD (tNAD, i.e. NAD+ plus NADH), a colorimetric assay (BioVision, Milpitas, California, USA, K337) was used on in mouse kidney tissue by colorimetric assays in accordance to the manufacturer’s protocol without usage of 10kd spin columns. These assays were performed on damaged and undamaged kidneys from the experiment illustrated in Fig. S4. Each sample was measured in duplicates in 1:10 or 1:20 dilution and for quantification an internal standard in duplicates was used. For normalization on protein amount, a standard BCA assay was used.

### Proteomic and transcriptomic raw data availability

Proteomic raw data are available at the ProteomeXchange Consortium via the PRIDE (63) partner repository (identifier: PXD024685; Username; Password: please ask). RNAseq data have been published before and are available at https://www.ebi.ac.uk/arrayexpress/experiments/E-MTAB-7982 or http://shiny.cecad.uni-koeln.de:3838/IRaP/.

### Study approval

The animal procedures were performed in accordance with the LANUV NRW (Landesamt für Natur, Umwelt und Verbraucherschutz Nordrhein-Westfalen, State Agency for Nature, Environment and Consumer Protection North Rhine-Westphalia; VSG 84-02.04.2013.A158 and 84-02.04.2017.A162).

Human samples were obtained as part of one clinical study at our center (64) (study name: CR_KCH, clinicaltrials.gov identifier: NCT01534364). The cohort consisted of patients that were provided with formula diet with an amount of 60% of daily energy expenditure (DEE) for 7 days prior to a coronary-arteria bypass graft. Blood samples included in our analyses were taken at study inclusion and after the diet after at least 5 h of fasting. The study was operated in accordance with the Declaration of Helsinki and the good clinical practice guidelines by the International Conference on Harmonization. All patients approved study inclusion and approval of each study protocols were obtained from the local institutional review board (Ethics committee of the University of Cologne, Cologne, Germany; protocol code 11-296).

## Author contributions

Experiments: MRS, KJRH-A, FCK, MJ, KK

Analyses of experiments: MRS, KJRH-A, MK, JUB, MH, SB, JWL, TK

Human sample acquisition: FG

Study design: TB, BS, VB, R-UM

Contribution of new tools: SB

Interpretation of results: MRS, KJRH-A, VB, R-UM

Writing – original draft: MRS, KJRH-A

Writing – review and editing: MJ, SB, KK, FCK, MH, JWL, TK, FG, TB, VB, R-UM

Method used in assigning the authorship order among co-first authors: Alternating order for first-co-authors, who collaborated in more than one project.

## Data and materials availability

All data are available in the main text or the supplementary materials.

## Acknowledgements

The authors acknowledge support from the CECAD Imaging Facility (head: Astrid Schauss), Dr. Monica Sanchez-Ruiz (Institute for Pathology, University Hospital of Cologne). Martyna Brütting (CECAD), Serena Greco-Torres (CECAD), Ruth Herzog (CECAD) and Stefanie Keller (CECAD) provided excellent technical support. We also acknowledge the support of the CECAD *in vivo* Research Facility and their Transgenic Core Unit for excellent mouse care and the generation of knockout mice (Branko Zevnik).

