## supplementary information for "Organ protection by caloric restriction depends on activation of the *de novo* NAD+ synthesis pathway"

### Supplementary figures

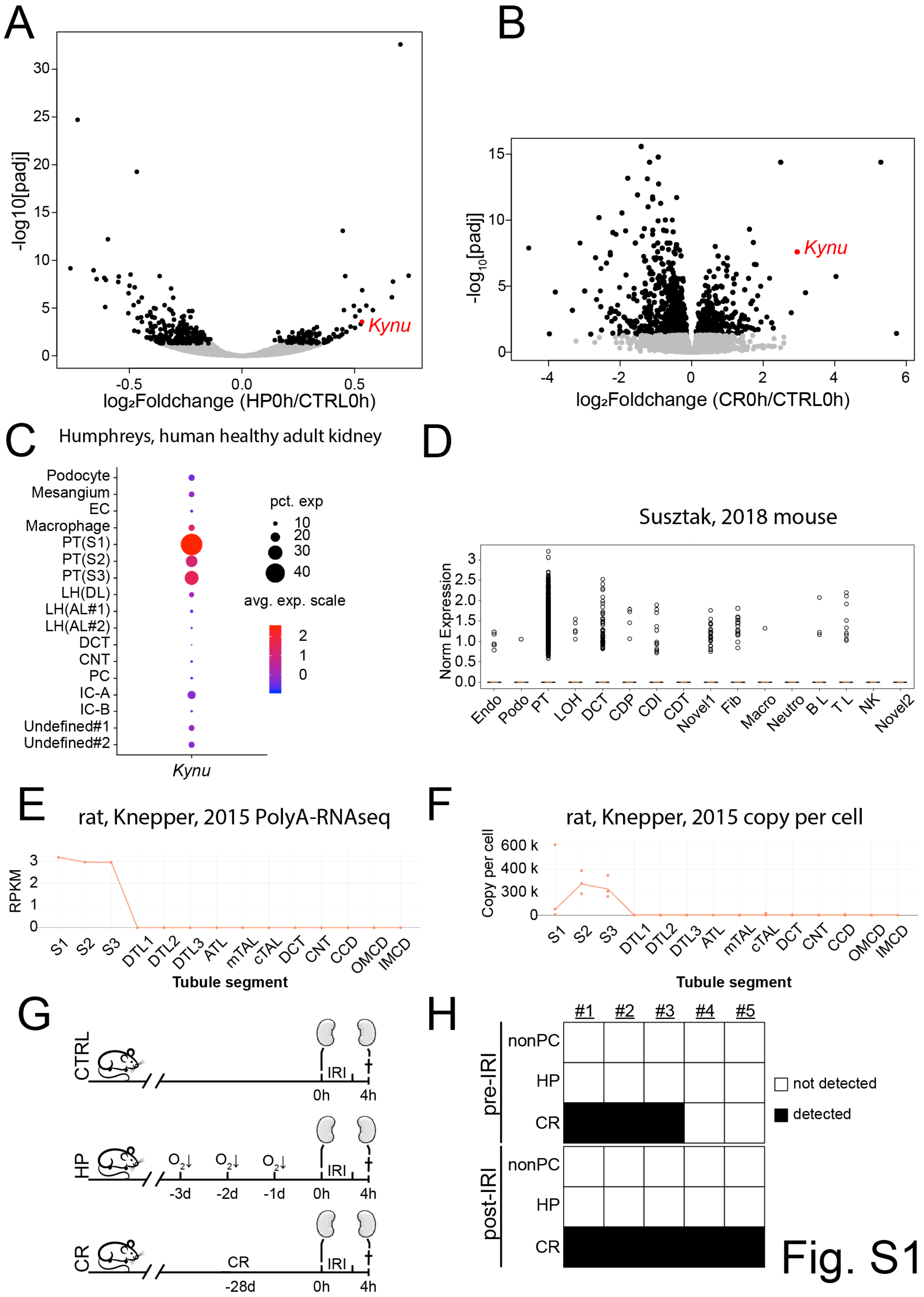


Fig. S1

Fig. S1: *Kynu* is differentially regulated by HP and CR and is mainly localized in the proximal tubulus.
**A, B.** Volcano plots showing the regulation of *Kynu* in murine kidneys after HP **(A)** or CR **(B)** compared to nonPC at 0 h prior to IRI. Gray, black and red dots depict genes without differential regulation (adjusted p>0.05), differentially regulated genes and *Kynu*, respectively. **C.** Bubble plot reflecting *Kynu* expression levels in a published single-cell RNAseq dataset of healthy human kidneys (Wu, Uchimura et al. 2018). **D.** Box plot chart reflecting means and interquartile range of *Kynu* expression levels in a published single-cell RNAseq dataset of adult murine kidneys (Park, Shrestha et al. 2018). **E.** Graph reflecting *Kynu* expression in microdissected tubuli segments of healthy adult rats (Lee, Chou et al. 2015). **F.** Graph reflecting KYNU levels in microdissected tubuli segments of healthy adult rats (Limbutara, Chou et al. 2020). **G.** Schematic illustration of the experimental setup to acquire kidneys for proteomic analyses. Mice were divided into three treatment groups (nonPC, HP, CR; n=5 per group) prior to IRI. Right nephrectomy was performed at the timepoint of surgery (0 h). To obtain blood and the left kidneys, 4 h after reperfusion all mice were sacrificed. Heatmap depicting KYNU detection of each analyzed sample by LC-MS/MS.
Abb.(alphabetic order): BL: B-lymphocytes; CD-PC: collecting duct-principal cell; CNT: connecting tubule; DCT: distal convoluted tubule; EC: endothelial cell; IC-A: intercalated cell type A; IC-B: intercalated cell type B; LH(AL): loop of Henle ascending loop; LH(DL): loop of Henle descending loop; MC: mesangial cell; PC: principal cell; Pod: podocyte; PT: proximal tubule; RPKM: Reads Per Kilobase Million; TL: T-lymphocytes; Endo: containing endothelial, vascular and descending loop of Henle; Podo: podocyte; PT: proximal tubule; LOH: ascending loop of Henle; DCT: distal convoluted tubule; CD-PC: collecting duct principal cell; CD-IC: CD intercalated cell; CD-Trans: CD transitional cell; Fib: fibroblast; Macro: macrophage; Neutro: neutrophil; NK: natural killer cell.

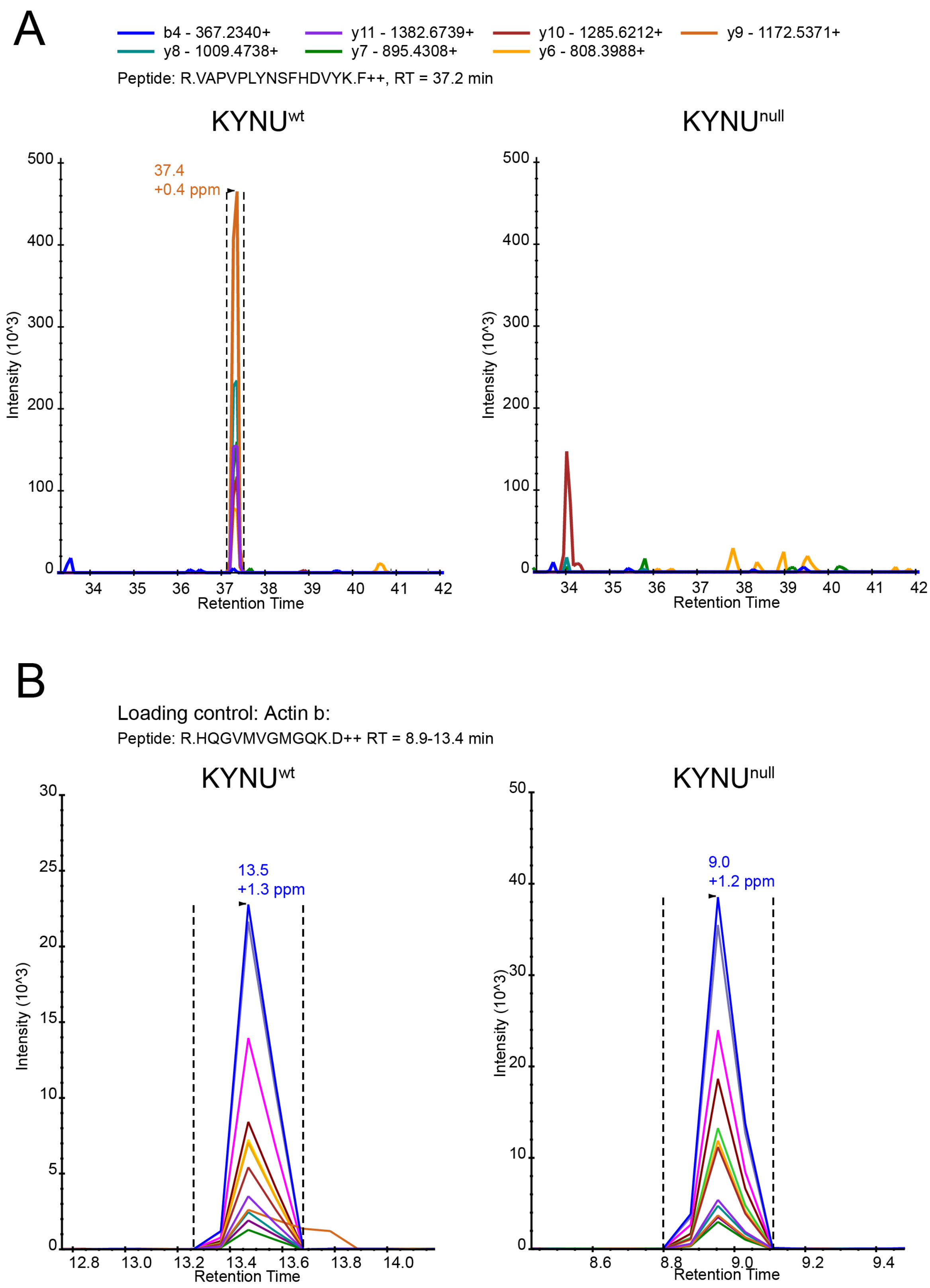
A
b4 - 367.2340+
y11 - 1382.6739+
y10 - 1285.6212+
y9 - 1172.5371+
y8 - 1009.4738+
y7 - 895.4308+
y6 - 808.3988+
Peptide: R.VAPVPLYNSFHDVYK.F++, RT = 37.2 min
KYNU^wt
KYNU^null
37.4
+0.4 ppm
Intensity (10^3)
Retention Time
B
Loading control: Actin b:
Peptide: R.HQGVMVGMGQK.D++ RT = 8.9-13.4 min
KYNU^wt
KYNU^null
13.5
+1.3 ppm
9.0
+1.2 ppm
Intensity (10^3)
Retention Time

Fig. S2

Fig. S2: Targeted proteomic analyses confirm KYNU-deficiency. Mass spectra of single kidney samples of KYNU wildtype *(*$KYNU^{WT}$) and KYNU deficient *(*$KYNU^{null}$) mice. **A.** Peptide R.VAPVPLYNSFHDVYK.F++, retention time (RT) = 37.2 min; **B.** Loading control: ACT b, Peptide R.HQGVMVGMGQK.D++, retention time (RT) = 8.9-13.4 min.

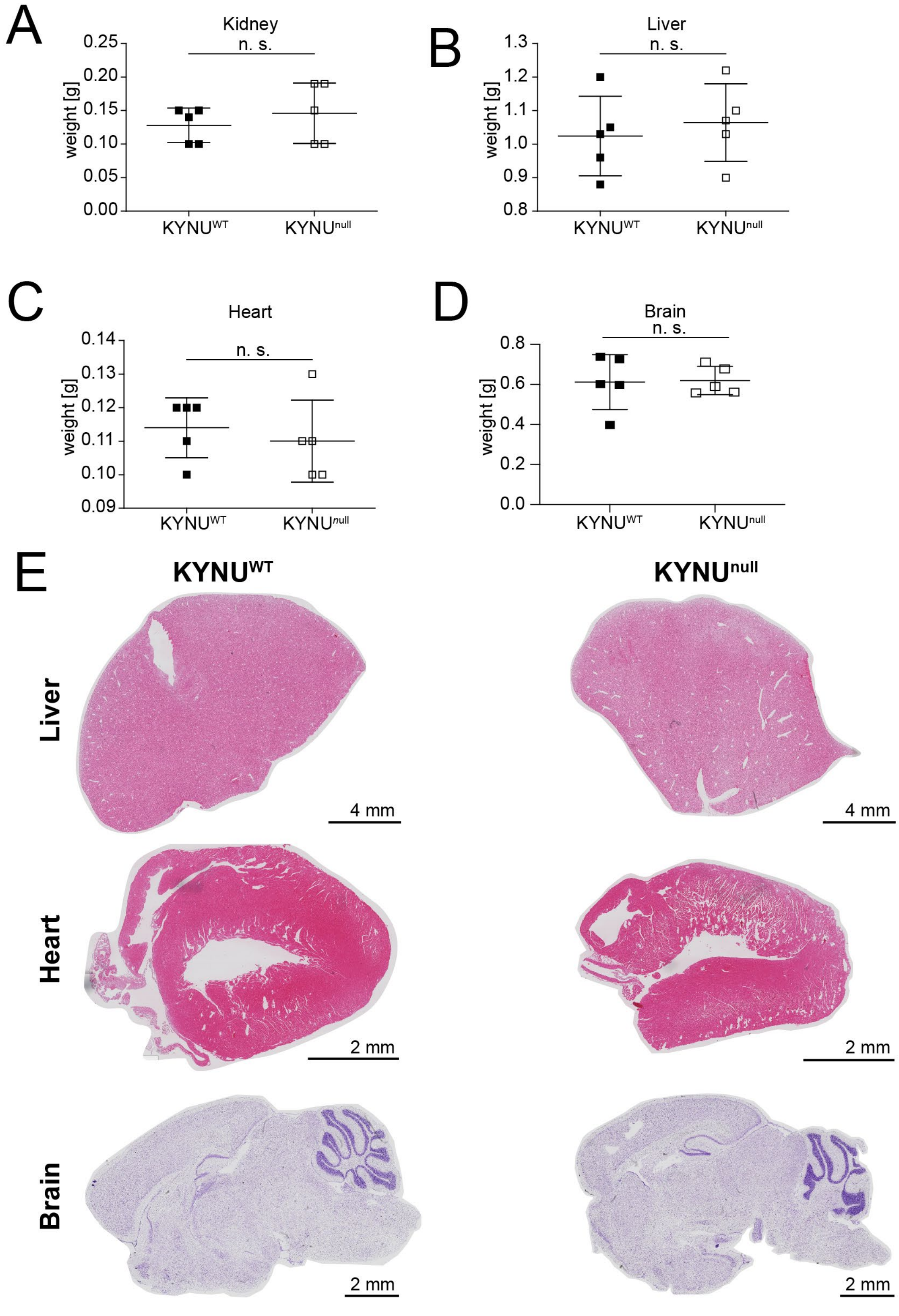


Fig. S3

Fig. S3: KYNU$^{null}$ mice do not show abnormalities concerning organ weights or basal histological analyses of the liver, heart or brain. **A.** Kidney weights of 10 to 14-week-old KYNU$^{WT}$ and KYNU$^{null}$ mice (unpaired student's t-test, n=5 per group). **B.** Liver weights of 10 to 14-week-old KYNU$^{WT}$ and KYNU$^{null}$ mice. **C.** Heart

weights of 10 to 14-week-old $KYNU^{WT}$ and $KYNU^{null}$ mice (unpaired student's t-test, n=5 per group). **D.** Brain weights of 10 to 14-week-old $KYNU^{WT}$ and $KYNU^{null}$ mice (unpaired student's t-test, n=5 per group). **E.** Hematoxylin-eosin stainings of livers and hearts as well as cresylviolett stainings of brains of 10 to 14-week-old $KYNU^{WT}$ and $KYNU^{null}$ mice.
**Abb.:** n.s.: p>0.05.

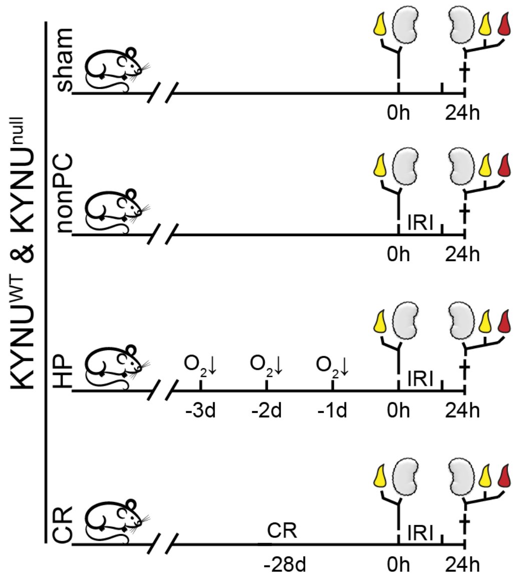


Fig. S4

Fig. S4: Experimental setup for phenotyping of KYNU$^{null}$ mice in the context of HP and CR and renal IRI. KYNU wildtype (KYNU$^{WT}$) mice and KYNU deficient (KYNU$^{null}$) littermates were each divided into 4 treatment groups (sham, nonPC, HP, CR; n=12 per group) prior to IRI. Right nephrectomy and urine collection (yellow drops) was performed at the timepoint of surgery (0 h). To obtain blood (red drops) and the left kidneys, 24 h after reperfusion all mice were sacrificed.
**Abb.:** sham: right nephrectomy followed by no-clamping of the left renal pedicle; nonPC: non-preconditioned male mice; HP: hypoxic preconditioning; CR: caloric restriction; d: day; IRI: ischemia-reperfusion injury.

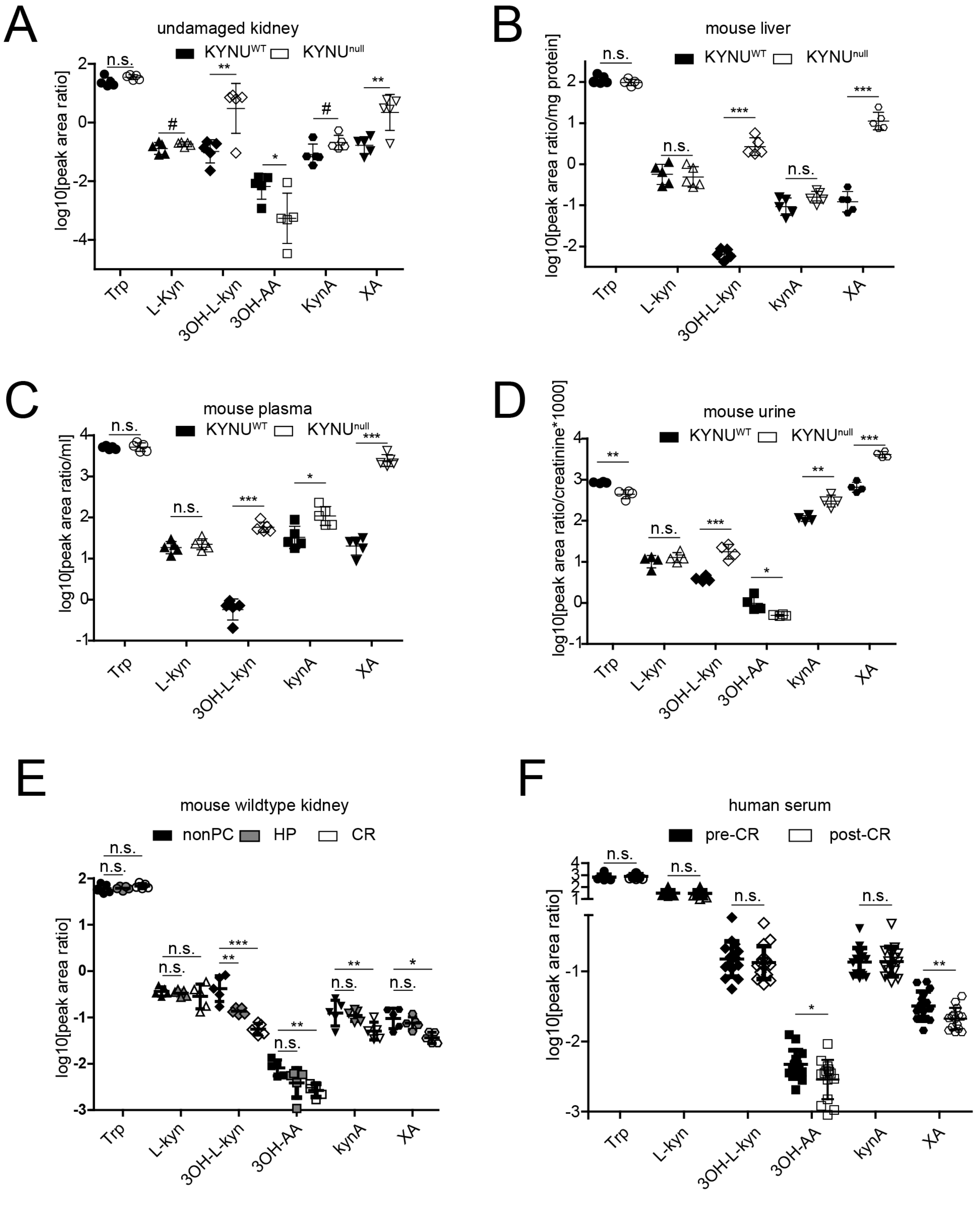
A
undamaged kidney
KYNUWT KYNUnull
log10[peak area ratio]
Trp
L-Kyn
3OH-L-kyn
3OH-AA
KynA
XA
n.s.
#
**
*
B
mouse liver
KYNUWT KYNUnull
log10[peak area ratio/mg protein]
Trp
L-kyn
3OH-L-kyn
kynA
XA
n.s.
***
C
mouse plasma
KYNUWT KYNUnull
log10[peak area ratio/ml]
Trp
L-kyn
3OH-L-kyn
kynA
XA
n.s.
***
*
D
mouse urine
KYNUWT KYNUnull
log10[peak area ratio/creatinine*1000]
Trp
L-kyn
3OH-L-kyn
3OH-AA
kynA
XA
**
n.s.
***
*
E
mouse wildtype kidney
nonPC HP CR
log10[peak area ratio]
Trp
L-kyn
3OH-L-kyn
3OH-AA
kynA
XA
n.s.
**
***
*
F
human serum
pre-CR post-CR
log10[peak area ratio]
Trp
L-kyn
3OH-L-kyn
3OH-AA
kynA
XA
n.s.
*
**

Fig. S5

Fig. S5: KYNU-deficiency and preconditioning affect metabolomic profiling of tryptophan metabolites. **A.** Quantification of tryptophan and its metabolites in undamaged kidney of $KYNU^{WT}$ and $KYNU^{null}$ mice (unpaired student's t-test, n=5 per group). **B.** Quantification of tryptophan and its metabolites in livers $KYNU^{WT}$ and $KYNU^{null}$ mice (unpaired student's t-test, n=5 per group). **C.** Quantification of tryptophan and its metabolites in blood plasma of $KYNU^{WT}$ and $KYNU^{null}$ mice (unpaired student's t-test, n=5 per group). **D.** Quantification of tryptophan and its metabolites in urine of $KYNU^{WT}$ and $KYNU^{null}$ mice (unpaired student's t-test, n=4 per group). **E.** Quantification of tryptophan and its metabolites in undamaged kidneys of $KYNU^{WT}$ and $KYNU^{null}$ mice without preconditioning (nonPC) or after HP/CR (one-way ANOVA and Tukey posthoc test, n=5 per group) **F.** Quantification of tryptophan and its metabolites in longitudinally acquired human blood serum of patients pre- and post-CR (paired student's t-test, n=15 per group).
n.s.: p-value>0.05; *: p-value< 0.01 to 0.05; **: p-value < 0.001 to 0.01; ***: p-value < 0.001

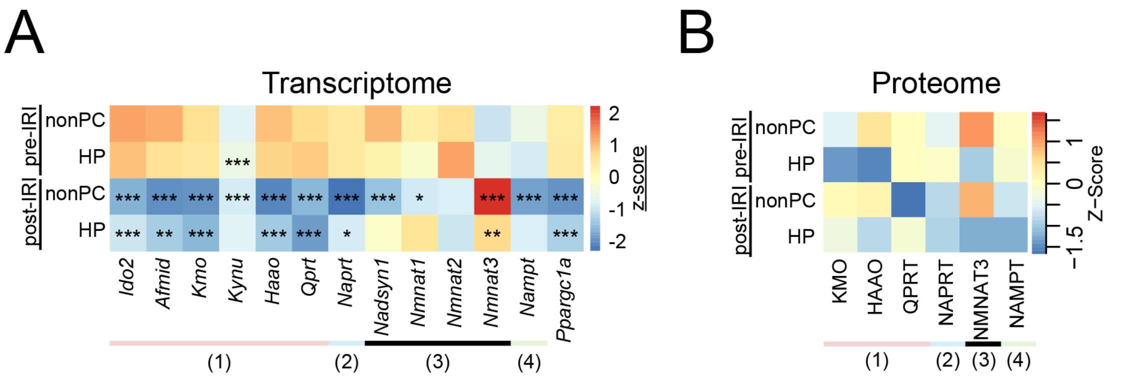


Fig. S6

Fig. S6: HP showed no influence on NAD levels and induced only weak transcriptomic or proteomic changes
**A.** Heatmap depicting mean based z-score expression levels of the genes involved in the NAD+ biosynthesis at 0 h pre-IRI and at 24 h post-IRI in non-preconditioned (non-PC) mice or after HP; asterisks reflect significance of pairwise comparisons of each gene between treatment groups (n=5 per group). **B.** Heatmap depicting median based z-score expression levels of the detected enzymes involved in the NAD+ biosynthesis at 0 h prior to (pre-IRI) and at 4 h after IRI (post-IRI) in non-preconditioned (nonPC) mice or after HP; asterisks reflect significance of pairwise comparisons of each gene between treatment groups (n=5 per group).
Abb.: (1): *de novo* branch of NAD+ biosynthesis; (2): Preiss-Handler branch of NAD+ biosynthesis; (3): belonging to 1, 2 and 4; (4): salvage branch of NAD+ biosynthesis. *: adjusted p-value/q-value<0.05, **: adjusted p-value/q-value: 0.001 – 0.01, ***: adjusted p-value/q-value<0.001

### Supplementary tables

| | | | | | | |
|---|---|---|---|---|---|---|
| | | | | CTRL0h | | |
| Ensembl ID | gene_symbo | C20 | C22 | C23 | C24 | HPC16 |
| ENSMUSG00( | Ido2 | 238 | 273 | 268 | 277 | 24 |
| ENSMUSG00( | Afmid | 452 | 485 | 707 | 882 | 211 |
| ENSMUSG00( | Kmo | 3103 | 3144 | 4043 | 4205 | 722 |
| ENSMUSG00( | Kynu | 107 | 95 | 126 | 114 | 70 |
| ENSMUSG00( | Haao | 3964 | 3524 | 4196 | 4715 | 367 |
| ENSMUSG00( | Qprt | 1532 | 1340 | 1587 | 1677 | 1007 |

Supplemental Table 1: Key enzymes involved in the three NAD+ biosynthesis pathways and their modulation by IRI and preconditioning. Raw read counts, normalized read counts, log2 fold changes and p-values for genes used for analyses of Fig. 6A are listed.

| Majority protein IDs | Protein names | Gene names (current) | meanLFQ(nPC_preIRI) | meanLFQ(nPC_post-IRI) | meanLFQ(CR_preIRI) | meanLFQ(CR_post-IRI) |
|---|---|---|---|---|---|---|
| Q78JT3 | 3-hydroxyan | Haao | 1.6531944 | 1.47892823 | 2.18321738 | 1.89486361 |
| Q8CC86 | Nicotinate p | Naprt | 1.35160379 | 1.3166358 | 1.52511761 | 1.59620419 |
| Q91WN4 | Kynurenine 3 | Kmo | 1.03955839 | 1.19913377 | 1.53938961 | 1.466242 |
| Q91X91 | Nicotinate-n | Qprt | 0.64944823 | 0.37251549 | 0.87060032 | 0.81855495 |
| Q99JR6 | Nicotinamid | Nmnat3 | -2.00731642 | -2.07866623 | -1.98081021 | -2.54371583 |
| Q99KQ4 | Nicotinamid | Nampt | 1.92392061 | 1.8812978 | 2.0355854 | 1.9941256 |

Supplemental Table 2: Changes on the proteome level in the *de novo* branch of NAD+ biosynthesis. Mean LFQ levels, log2 fold changes and statistical analyses for the data of Fig. 6B are listed.

| Q1 Mass (Da) | Q3 Mass (Da) | ID | DP (volts) | EP (volts) | CE (volts) | CXP (volts) |
|---|---|---|---|---|---|---|
| 114.0 | 44.0 | Creatinine | 70 | 10 | 46 | 10 |
| 117.0 | 47.0 | D3 Creatinine | 70 | 10 | 46 | 10 |
| 154.1 | 80.0 | 3OH Anthranili | 11 | 10 | 33 | 10 |
| 154.1 | 108.0 | 3OH Anthranili | 11 | 10 | 27 | 12 |
| 154.1 | 136.0 | 3OH Anthranili | 11 | 10 | 15 | 14 |
| 190.0 | 89.0 | Kynurenic acic | 46 | 10 | 50 | 12 |
| 190.0 | 116.0 | Kynurenic acic | 46 | 10 | 41 | 14 |
| 190.0 | 144.1 | Kynurenic acic | 46 | 10 | 25 | 16 |
| 195.1 | 94.0 | D5 Kynurenic a | 51 | 10 | 50 | 12 |

Supplemental Table 3: MRM transitions and compound-specific parameters. Metabolites were monitored in the positive ion mode with their specific Multiple Reaction Monitoring (MRM) transitions. Values for declustering potential (DP), entrance potential (EP), collision energy (CE) and cell exit potential (CXP) of the different MRM transitions are listed.
